# Chance favors the prepared genomes: horizontal transfer shapes the emergence of antibiotic resistance mutations in core genes

**DOI:** 10.1101/2023.06.20.545734

**Authors:** Charles Coluzzi, Martin Guillemet, Fanny Mazzamurro, Marie Touchon, Maxime Godfroid, Guillaume Achaz, Philippe Glaser, Eduardo PC Rocha

**Affiliations:** Institut Pasteur, Université Paris Cité, CNRS, UMR3525, Microbial Evolutionary Genomics, Paris, France; Collège Doctoral, Sorbonne Université, Paris, France; SMILE group, Center for Interdisciplinary Research in Biology (CIRB), Collège de France, CNRS, INSERM, Université PSL, Paris, France; Institut Pasteur, Université de Paris Cité, CNRS, UMR6047, Unité EERA, Paris, 75015, France

## Abstract

Bacterial lineages vary in the frequency with which they acquire novel traits, like antibiotic resistance or virulence. While previous studies have highlighted the impact of the genetic background on the successful acquisition of novel traits through horizontal gene transfer, the impact of the latter on the subsequent evolution of bacterial genomes by point mutations remains poorly understood. Here, we studied the evolution of resistance to quinolones in thousands of *Escherichia coli* genomes. Resistance-conferring point mutations in the core genes are frequent and accumulate very quickly. We searched for gene gains and losses significantly associated with the subsequent acquisition of these resistance mutations. This revealed 60 groups of genes in genetic linkage whose gain or loss induced a change in the probability of subsequently becoming resistant to quinolones by point mutations in *gyrA* and *parC*. Although some of these chronologies may reflect epidemiological trends, most of these groups encoded functions that were previously associated with antibiotic resistance, tolerance, or persistence, often specifically under quinolone treatment. A lot of the largest groups were found in prophages or plasmids, and they usually increased the likelihood of subsequent resistance mutations. Conversely groups of lost genes were typically small and chromosomal. Quinolone resistance was among the first resistances acquired in the extant lineages of *E. coli* and its acquisition was associated with an increased likelihood of acquiring other types of resistances, including to aminoglycosides and beta-lactams. Our findings suggest that gene flow shapes the subsequent fixation rate of adaptive mutations in core genes. Given the substantial gene flow within bacterial genomes, interactions between horizontal transfer and point mutations in core genes may be key to the success of adaptation processes.

## Introduction

Bacterial populations adapt rapidly to novel challenges such as bacteriophage predation, antibiotic therapy, or environmental perturbations. Adaptation is facilitated by point mutations and by genetic exchanges with other bacteria. Since the genome is made of thousands of genes linked in complex regulatory networks and encoding proteins with multiple physical interactions, these modifications may impact other processes than those directly involved in adaptation. The trade-off between the benefits and the cost of the novel variants shapes the outcome of their natural selection. Several studies have shown how the acquisition of novel functions by horizontal gene transfer (HGT) depends on the existing genetic background (1, 2). For example, metabolic pathways grow by transfer of genes encoding enzymes involved in reactions at the periphery of existing networks (3), whereas genome reduction involves an ordered loss of specific functions (4–6). Interactions between the genetic background and genes acquired horizontally may tune the probability of fixation of the latter. Such interactions may be key to understand the adaptation of bacteria because their gene repertoires vary rapidly by HGT. For example, less than half of the average *Escherichia coli* genome corresponds to genes present in more than 99% of the strains (core genome). Consequently, the diversity of *E. coli* gene repertoires, its pan-genome, is more than an order of magnitude larger than the average genome (7, 8). The large genomic variation caused by HGT and gene loss means that genetic backgrounds can be very different within a species. Hence, not only HGT may be affected by the genetic background, it’s likely that the evolutionary trajectories of core genes are also affected by changes in gene repertoires. There are very few studies on the existence of the epistatic interactions between HGT and point mutations.

The evolution towards antibiotic resistance in bacterial pathogens (and commensals) is a recent example of the ability of bacteria to rapidly adapt to novel challenges (9). Bacteria can counteract antibiotic therapies by diminishing the intracellular concentration of the antibiotic by reducing its influx or increasing its efflux. They can also protect the target of the antibiotic or modify it by mutation. Finally, they can inactivate the antibiotic using appropriate enzymes or evolve alternative pathways that make the target redundant (target bypass) (10, 11). These mechanisms may be acquired by mutation or by HGT (12). While high environmental pressure imposed by antibiotics is central to the emergence of antibiotic resistance (13, 14) epistatic interactions were shown to shape it (15–19). The evolutionary trajectories towards resistance are constrained by the existence of favorable mutational paths where intermediate steps have lower-than-average fitness costs and/or higher than average resistance (20–23). Epistatic interactions may result in the fitness cost of a resistance being alleviated by the presence of another one, which may favor the evolution of multi-drug resistance (24–27). For instance, the high transmission fitness of multiple drug resistance *Mycobacterium tuberculosis* strains of lineage 2 resulted from epistatic interactions between compensatory mutations in RNA polymerase and the rifampicin resistance-conferring mutation RpoB S450L (28). On the other hand, collateral sensitivity occurs when resistance to an antibiotic is linked with an increased susceptibility to another antibiotic (10, 29).

Differences in the genetic background influence the evolution of antibiotic resistance (30–32), even under strong selection (33). However, few studies identified chronologies between specific changes in the genetic background and the acquisition of the antibiotic resistance (34). Here we define chronologies as a set of events ordered in time. In a seminal study, the fitness cost of chromosomal resistance to several antibiotics acquired by point mutations was found to be in negative epistatic association with the presence of conjugative plasmids in more than half of the tested combinations (35). More recently, the presence of the efflux pump *norA* potentiated the subsequent evolution of point mutation conferring resistance to quinolones in *S. aureus* (36). Finally, the ability of *Escherichia coli* to evolve high-level colistin resistance by point mutation in the core gene *lpxC* increased in the presence of a plasmid carrying the *mcr*-1 colistin resistance gene (37). Epidemiological data seems to confirm these trends. For example, the *E. coli* strains from the clone ST131 comprise now the majority of extended spectrum beta-lactamase (ESBL)-producing isolates from the species (acquired by HGT) and they are almost invariably also resistant to fluoroquinolones (acquired by point mutations)

(38). Hence, interactions between genes acquired by HGT and chromosomal mutations may be important. Given the differences in terms of genetic backgrounds across a bacterial species, caused by rampant HGT, epistatic interactions may contribute to explain why strains from certain lineages are more often antibiotic resistant than others (39, 40).

We study the evolution of resistance to quinolones, a large group of broad-spectrum bactericidals widely used in human and veterinary medicine (41–43). In 2017, they represented 9.5% of the antibiotic usage in the 30 European union countries (44). Quinolones disrupt the function of the bacterial type II topoisomerases by inhibiting the catalytic activity of DNA gyrase (*gyrA* and *gyrB*) and topoisomerase IV (*parC* and *parE*). DNA gyrase introduces supercoils and DNA topoisomerase IV prevents supercoiling from reaching unacceptably high levels. These proteins intervene during replication or transcription and their mechanism involves the creation of a double strand break that is subsequently ligated. Quinolones stabilize the cleavage complex preventing the enzymes from performing the ligation step, thereby resulting in double strand breaks that may lead to replication stalling and cell death (45–47). The primary target for quinolones is the gyrase in enterobacteria and the topoisomerase in Firmicutes (45). The mechanisms that provide high resistance to quinolones have been characterized in detail. They involve mutations in quinolone-resistance determining regions (QRDR) of the target proteins. In *E. coli*, mutations appear most frequently at codons 83 and 87 of *gyrA*, near the active site of the gyrase, altering the binding of quinolones and reducing the cell’s susceptibility to them (48). Mutations in *gyrB* are rarer and some provide higher susceptibility to specific quinolones, a trait that can be masked by epistatic interactions with *gyrA* mutations (49). Mutations in the topoisomerase subunits *parC* and *parE* of *E. coli* usually co-occur with mutations in *gyrA* (50, 51), suggesting that the mutations in topoisomerase will not be fixed unless the sensitivity of the DNA gyrase has been reduced. Interestingly, a *parC* mutation was shown to strongly alleviate the cost of *gyrA* mutations (22). The relevance of epistatic interactions in the target protein mutations leading to quinolone resistance has been shown in many species, *e.g. E. coli* (52), *Streptococcus pneumoniae* (53), *Pseudomonas aeruginosa* (54) and *Mycobacterium tuberculosis* (32). A combination of sequence analysis of 195 resistant clinical isolates, experimental work and mathematical modeling revealed the major trajectories of ciprofloxacin-resistance evolution in *E. coli*, explaining the prevalence of a few dominant genotypes and the order of accumulation of the mutations (55). Epistatic interactions with other determinants of resistance for other antibiotics have also been reported (56–58) and shown to be sensitive to changes in the genetic background (59). Of note, lower resistance-level to quinolones can also be provided by plasmid-borne genes such as *qnrA*, *qepA*, AAC(6′)-Ib and *oqxAB* (60–63)

Here, we wished to know if there is evidence in natural populations that gene gains and losses change the propensity of bacteria to adapt by specific point mutations that are known to result in antibiotic resistance. For this, we identified events of gene gain and loss in the *E. coli* phylogenetic tree that occurred systematically in branches preceding those where resistance to quinolones took place. This approach allows to explicitly account for population structure when counting events. We then computed the *induction* effect (termed here lambda) that one event has on the other, i.e., if a given gene gain or loss increased or decreased the frequency of subsequent acquisition of antibiotic resistance by point mutations. Of note, this approach is different from the ones used in bacterial GWAS analyses where the goal is to identify frequent co-occurrences of events in the same branch or simultaneous presence of genes in extant taxa (64–67). Here, the goal is to identify changes that were fixed in the lineage before the acquisition of mutations conferring resistance to quinolones. We focus on the latter because they are well known and occur in core essential genes. Also, it was shown that the same point mutation in *gyrA* conferring resistance to quinolones can have very different costs depending on the genetic background (68), a clear indication that pre-existing variation in the latter may shape evolution in core genes.

## Results

### Frequency of the mutations providing resistance to quinolones

We analyzed 1 268 complete genomes of *E. coli* from RefSeq and 877 draft genomes of isolates sampled across multiple sources in Australia (named AUS, (69)). The first is our reference dataset and the second is mentioned occasionally because it has much fewer clinical isolates (but the genomes are not fully assembled). The 2 145 genomes have 78 653 different gene families (pangenome) and 2 393 core gene families (genes present in a single copy in at least 99% of the genomes). The latter were used to construct a phylogenetic tree for each dataset (Figure 1, S1). We built the trees using the same set of core genes to have comparable phylogenies and phylogeny-based analyses in the two datasets. The trees were well-resolved with more than 85% of the branches having more than 90% Ultra Fast Bootstrap.

**Figure 1:**
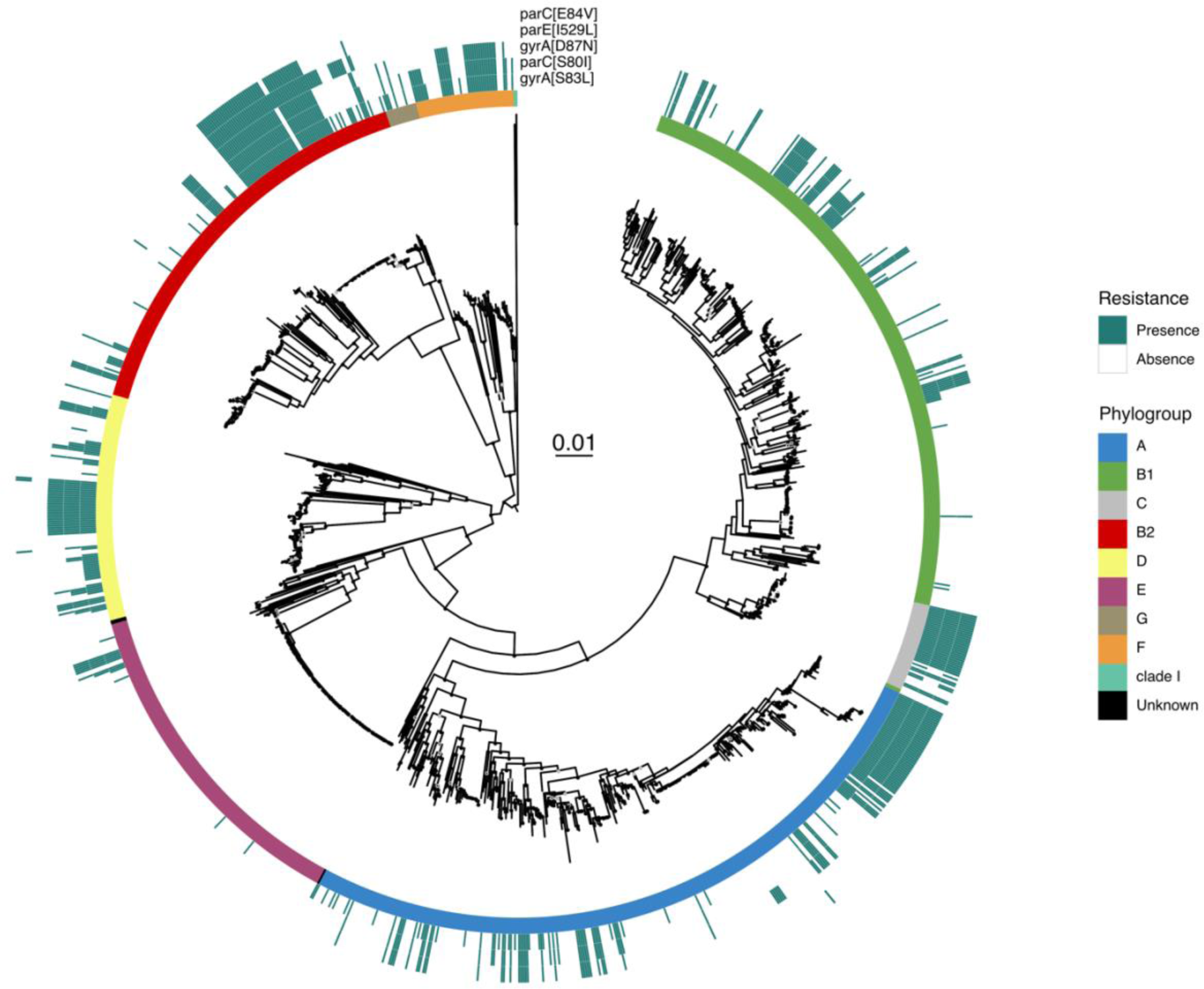
Phylogeny and distribution of quinolone resistance mutations in DNA gyrase (*gyrA*) and topoisomerase IV (*parE* and *parC*) genes in *E. coli* (RefSeq dataset). The maximum likelihood phylogenetic tree of the species was built with IQTree (model GTR+F+I+G4 and 1000 ultrafast bootstraps) the scale of substitution per site is in the center. The tree was midpoint rooted. The *E. coli* phylogroups are represented by the colors in the inner center. The marine green squares indicate the presence of the most frequent mutations across the species. Ultrafast bootstrap values superior to 75% are shown with a light gray circle and values superior to 90% with a black circle (see supplementary files).

We extracted from the proteome of the RefSeq dataset the sequences of the *gyrA, gyrB, parC* and *parE* genes. We screened these sequences for 39 different quinolone resistance mutations retrieved from the literature (Table S1). The mutations were remarkably frequent in the RefSeq dataset, where 39% of the strains had at least one mutation (Table S2). As a comparison, the AUS dataset had fewer resistance mutations (11% of the strains, Table S3) and these were concentrated in the human and poultry isolates (86.4%, Table S4). Hence, the absolute frequency of these mutations in the RefSeq dataset should be interpreted in the light of the presumed high frequency of clinical strains in the set. The search for antibiotic resistance genes (ARG) in the RefSeq dataset showed that resistance to quinolones (including the point mutations, *qnr*/*qep* and the plasmid-encoded multidrug efflux pump *oqx*) is the most frequent in the species, since 637 strains out of 1268 have at least one of the three types of resistance (Figure 2A, Table S5). Among these mechanisms of resistance to quinolones, the ones based on mutations were by far the most frequent.

**Figure 2:**
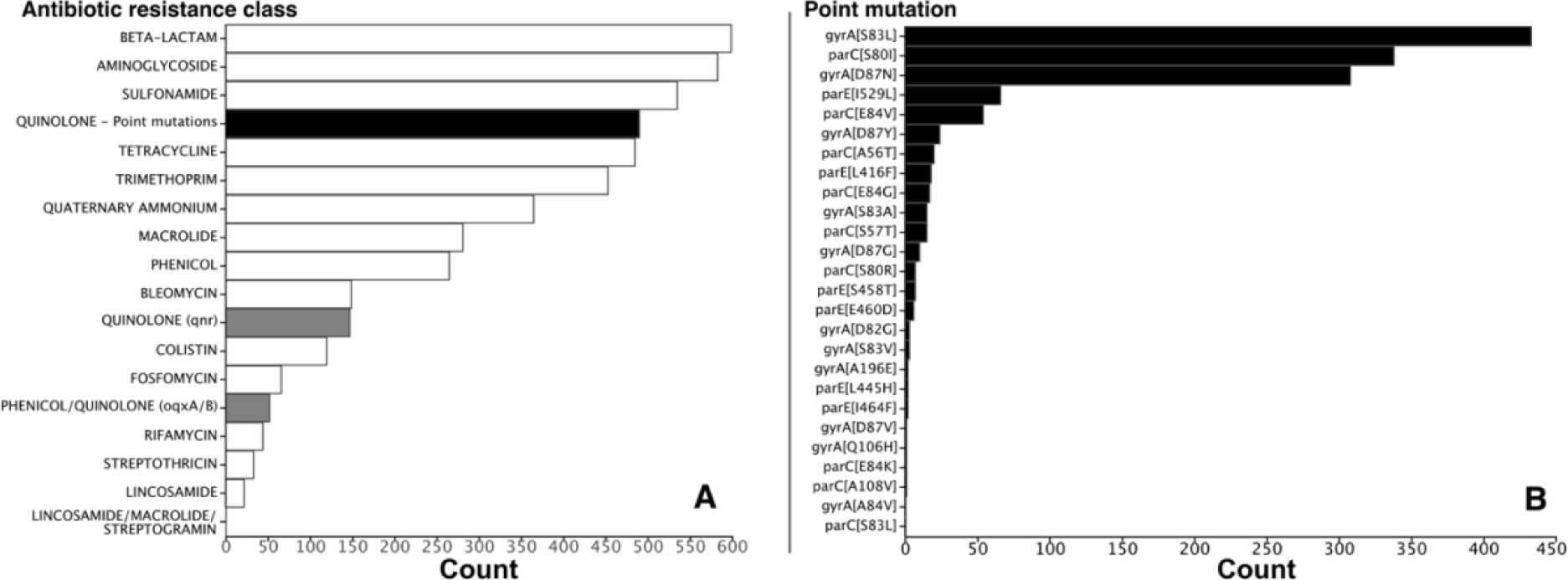
Distribution of the antibiotic resistance classes detected in the RefSeq dataset. **A:** Antibiotic resistance classes are ordered according to their frequency (total number in the dataset). Resistances to quinolone are separated into 3 distinct classes: resistance conferred by point mutations of core genes (black), by acquisition of *qnr* genes or *oqx* genes (grey). **B:** Mutations are ordered according to their frequency. The first letter in the name in the square brackets correspond to the ancestral amino acid, the number to the position of the transition and the last letter to the amino acid conferring the resistance, e.g., *gyrA*[S83L] means that the mutation is in *gyrA* at position 83 and involves a substitution Ser->Leu.

The mutations conferring resistance to quinolones had very different prevalence in the species (Figure 2B). The single mutation *gyrA*[S83L] which confers a significant increase in resistance at a low cost (70) is present in 34% of the strains. The two other most frequent mutations (*parC*[S80I] and *gyrA*[D87N]) were both present in more than 26% and 24% of the strains, respectively. The list of mutations in order of frequency then includes a mutation in *parE* and another in *parC*. Of note, mutations in *parE* and *gyrB* are much less frequent than those in *gyrA* and *parC*. Apart from the sheer abundance of mutations, higher in the RefSeq than in the AUS genomes, both datasets showed similar patterns in terms of the most frequent types of mutations (Figure S2). Hence, and in agreement with the literature (71, 72), a group of 3 mutations in two (of the four) key genes targeted by quinolones (*gyrA*[S83L], *gyrA*[D87N], *parC*[S80I]) are by far the most frequent in *E. coli*.

We then mapped the resistant isolates on the phylogenetic tree. We found them scattered across the species. They are present in all phylogroups, but their prevalence can be quite different. In phylogroup E and B1 these mutations are rare, whereas in phylogroup F they are very frequent. Hence, there is a non-uniform distribution of mutations for resistance across the species, with certain clades containing many more mutations than others (Figure 1). The AUS dataset shows similar results, with high frequency of resistance mutations in phylogroup F and low in E and B1 (Figure S1). Overall, the clustered distribution of resistances and the frequent presence of multiple mutations suggest that they accumulate in non-random ways.

### Order of acquisition of the mutations conferring resistance to quinolone

The phylogenetic analysis shows a strong co-occurrence of the different mutations conferring quinolone resistance. It is well-known that the combination of *gyrA*[S83L], *parC*[S80I], and *gyrA*[D87N] confers high level of resistance with limited fitness cost (22, 73). These three mutations co-occurred much more frequently together (182) than separately (90). The other low frequency mutations are often associated with them (Figure S3). Only one relatively rare mutation (*gyrA*[S83A]) was more typically found alone than in combination with the three major ones (Table S2, Figure S3). The systematic identification of large combinations of mutations raises the possibility that there exist typical chronologies for their accumulation. Indeed, it was shown using a combination of experimental biology and modeling that given the resistance they provide, the most likely path to adaptation was *gyrA*[S83L] -> *parC*[S80I] -> *gyrA*[D87N] (55).

To test this hypothesis, we searched for the preferential chronologies of the five main mutations on the RefSeq data using the species phylogenetic tree (see Methods, Figure 3). We did not consider the reverse mutations because these are very rare (1.7% of all paths across the tree), and because reversions are sometimes associated with poorly resolved regions of the tree and may thus be spurious. We observed similar trajectories towards the acquisition of the 3 main mutations in AUS dataset (Figure 3 and S4, Table S6, S7). In the RefSeq data, we identified 48 occurrences of lineages acquiring only the *gyrA*[S83L] mutation. This suggests that the mutation *gyrA*[S83L] is the first one fixed in most lineages. Some of the clones with this mutation then gave rise to the triple mutated strain. The other most frequent trajectory was the one going from the fully sensitive combination to the triple mutant (37 occurrences). Hence, our analysis failed to reveal unequivocally the substitution paths from the ancestral sensitive state to the triple mutant (*gyrA*[S83L], *parC*[S80I] and *gyrA*[D87N]). This suggests that the other two major mutations are acquired and fixed very quickly, such that they are inferred to arrive jointly in a single branch of the phylogenetic tree.

**Figure 3:**
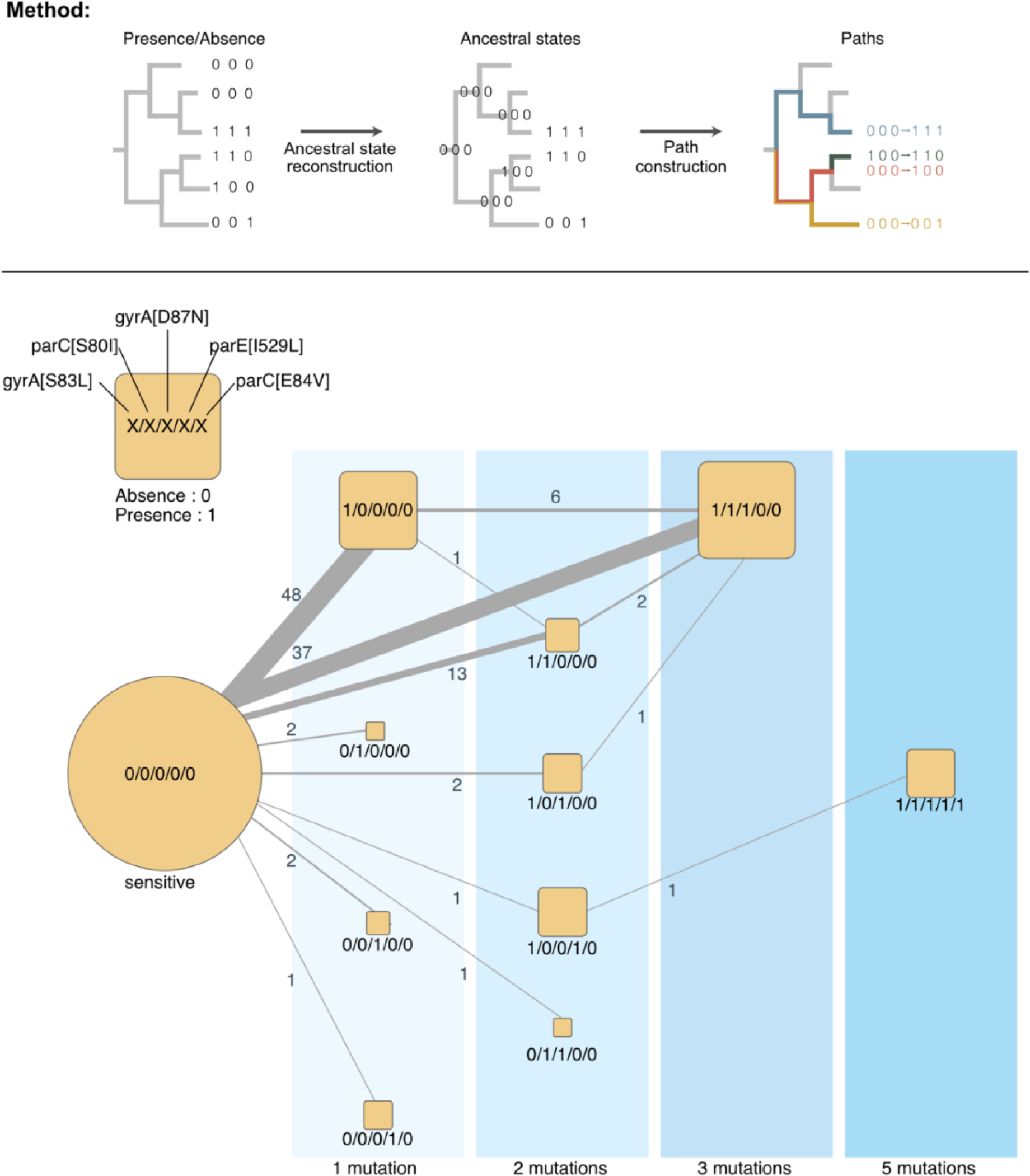
Chronologies of acquisition of the main resistance mutations in *E. coli* of the RefSeq dataset. The blue areas represent the number of distinct mutations in genomes. The size of the square scales with the number of genomes observed (leaves of the tree). The edges represent the chronologies of acquisition of one or several mutations as inferred from the reconstruction of ancestral states on the tree. The edge size is proportional to the frequency of the respective transition, with the labels showing this exact number.

To identify the paths towards the triple mutant, we used three genomic datasets already available in our laboratory consisting of very closely related genomes (one single ST): 757 of ST410, 85 ST131 and 579 for ST167. In ST410, out of the 757 strains, most (656) contain mutations conferring resistance to quinolone. This trait is not ancestral, as shown by the phylogenetic reconstruction of the sequences (Figure S5). Yet, almost all the resistant clones (653/656) harbor the *gyrA*[D87N]/*gyrA*[S83L]/*parC*[S80I] combination of mutations and only 2 among the basal isolates contain the *gyrA*[S83L] alone. We found similarly inconclusive results for the other STs. Hence, even at this fine level of granularity, one cannot reconstruct the precise order of fixation of the three mutations (Figure S6, S7, S8, S9 and S10, Table S8).

The quick accumulation of the three mutations could be due to homologous recombination replacing in one single event the ancestral sequences by the triple mutant. This seems unlikely because *parC* and *gyrA* are 1.39 Mb apart in the strain MG1655 and even *parC* and *parE* are more than 8 kb part (while observed average recombination tracts in *E. coli* are less than 1kb (74)). Still, we searched for evidence of recombination in the genomes using Gubbins. This analysis showed that in 7 out of the 39 events of acquisition of the two mutations in *gyrA* there was a recombination tract that was acquired overlapping the position of the mutations at the same time (same branch in the tree). This suggests that recombination may occasionally contribute to the emergence of the double mutant in the lineages. Yet, out of the 37 events of acquisition of the 3 mutations in a branch of the tree, only one coincided with the acquisition of recombination tracts at both genes (different tracts). Overall, these results suggest that *gyrA*[S83L] is the main initial driver of the evolution of quinolone resistance in *E. coli*, and that the subsequent mutations are fixed almost simultaneously by rapid accumulation of novel point mutations, although recombination may occasionally accelerate the process.

### Clusters of gene gains and losses shaping the emergence of resistance mutations

To investigate how the dynamics of gene repertoires favor the acquisition of resistance to quinolones by point mutations in the core genes targeted by the antibiotics, we searched for genes that were frequently gained or lost before the emergence of these mutations. Since most resistance mutations co-occurred, and they were all shown to have significant individual effects on resistance, we defined taxa as resistant when they had at least one resistance mutation. We then reconstructed the ancestral states of resistance (as defined above) and of each gene family of the pan-genome to infer the moments of gains and losses of the gene families and of the resistance. Finally, we used Evo-Scope to compare the chronology of gains and losses of every family in the pangenome with the acquisition of the resistance to the antibiotic. Of note, this software searches for successions of events in different branches of the tree. The two events are thus well-separated in time. This allowed to identify the significant chronologies, i.e., when one event of gene gain or loss was followed by the acquisition of resistance more frequently than expected given the frequency of events and their distribution on the phylogenetic tree (75). We identified 183 gene gains and 26 gene losses occurring frequently before the acquisition of the resistance (p<10^-5^ after correction for multiple tests, Figure 4, Table S9).

**Figure 4:**
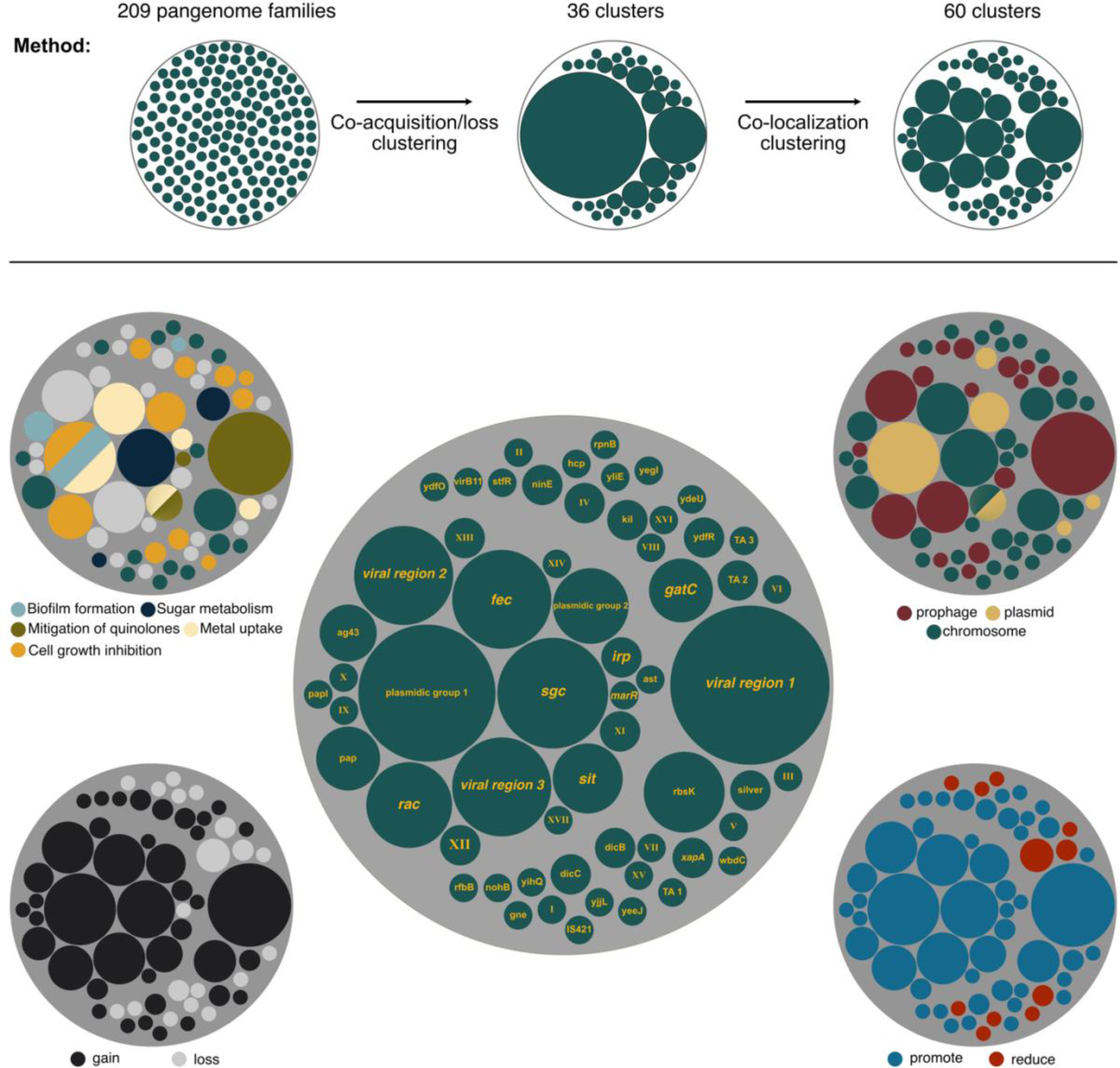
Cluster of genes consistently acquired or lost prior the acquisition of the quinolone resistance. The center figure represents the different clusters according to their size. Clusters with known functions or encoded by a specific mobile genetic element type were named accordingly. Others were given Roman numbers. Clusters are represented according to their location on prophage, plasmid or chromosome (top right), to their influence on the acquisition of the quinolone resistance (bottom right), to the nature of the event (bottom left) and to the functions they encode (top left).

Our method searches for preferential chronologies between a type of event that occurs in a set of branches of the phylogenetic tree and resistance mutations that emerge later (on higher branches). It is not aimed at identifying events that are associated, such as genes directly involved in the phenotype. Yet, it may also reveal associations in the sense usually employed in genome-wide association studies (GWAS). To assess if chronologies differ significantly from GWAS, we performed a GWAS analysis to identify genes whose presence or absence is associated with quinolone resistance. We then identified those positively or negatively associated with the trait (quinolone resistance) while accounting for the phylogenetic structure of the data using a linear mixed model (see Methods). This analysis yielded no significant associations at the same statistical threshold (correction for multiple tests and p<10^-5^). Lowering the significance threshold revealed 53 genes (P<0.01). Most of the 20 genes positively associated with quinolone resistance encode transporters, regulators, transposases, or enzymes associated with antibiotic resistance (*e.g.*, beta-lactams or tetracycline, Table S10). None of the 209 genes involved in the chronologies identified by the Evo-Scope analysis was identified by the GWAS analysis. This suggests that our chronologies identify events that favor the acquisition of resistance without contributing very strongly to it. This is not very surprising, considering that strong resistance is well-known to be due to mutations in a few core genes (and that was how it was defined in the GWAS).

Many of the events identified in the chronologies were acquisitions (see below, Figure 4). Genes acquired by HGT are often initially grouped in large mobile genetic elements or at least in fragments of DNA with more than one gene. If one of these genes is biologically associated with the emergence of resistance, other genes under strong genetic linkage, e.g., entering the genome in the same event of transfer, may also be identified as significantly associated with the emergence of resistance. To control for the effect of genetic linkage, we clustered the genes in function of their tendency to be acquired or lost in the same branches of the species tree (Table S11, S12, S13). We then sub-divided these clusters using information on genomic co-localization to group genes that tend to be neighbors (see Methods, Figure 4, Table S14). This led to the identification of 26 clusters of genes and 34 isolated genes (singletons) (Figure 4, Table S15). Hence, our analysis suggests that there is a minimal number of 60 genes whose gain/loss affect the probability of acquisition of resistance mutations.

The Evo-Scope method provides a parameter (λ) which is a measure of the impact of an event (here gene gain or loss) in increasing the likelihood of another (acquisition of resistance mutations) in a higher branch of the tree. These values were not used for the clustering but are expected to be similar among genes under strong genetic linkage. Indeed, genes in clusters had homogeneous lambda values relative to the others (levene Test: 5.71e-06, MSE (mean square of the error) =0.591, one-way ANOVA ***)), meaning that genes within the same cluster have approximately similar impacts on the acquisition rate of the quinolone resistance from the statistical point of view. Out of the 60 clusters, the majority (49) promotes the acquisition of the resistance mutations (λ>1), while only 11 clusters reduce it (λ<1) (Figure 4, bottom right). Most of the latter are close to the significance threshold. This means that most of the significant events of gene gain and loss associated with mutations tend to increase the likelihood of the resistance mutations.

The clustering procedure aimed at identifying blocks of genes that were acquired at the same time (along the same branch). As expected, all the clusters were exclusively made of groups of either lost or gained genes, not both. Out of the 60 clusters, the majority (41) correspond to gene gains and 19 correspond to gene losses. One might expect that the first correspond to gain of functions and the latter to losses. Bacteria need to acquire all the genes involved in a function to express it. However, the loss of only few genes is sufficient to disrupt the activity of entire pathways. This may explain why our analysis reveals that groups of gains are larger than groups of losses (Mann–Whitney Test: pvalue=0.0108) (Figure 4, bottom left). The acquisition of functions in large mobile genetic elements (MGE) may also contribute to explain these results. Except for one cluster that was either part of a plasmid or of a chromosome (depending on the genome), clusters were made exclusively of either plasmid, prophage genes or neither of the two. Eighteen clusters, including the largest, were in prophages. All the prophage clusters are associated with promotion of resistance mutations. Five clusters were on plasmids including the second largest (Figure 4, top right). In conclusion, our analysis identified 60 groups of genes, most of them corresponding to acquisitions and some of which in MGEs, that tend to increase the likelihood of acquisition of resistance mutations.

### Acquired functions that induced subsequent resistance acquisition

Among the 60 clusters identified above, 17 lack genes with predicted functions. The other clusters tend to have many genes involved in closely related functions, organized in loci with the characteristics of operons (co-oriented closely spaced genes). The analysis of the AUS dataset revealed much fewer genes (62) significantly associated with the emergence of quinolones (Table S16), which is probably caused by the much lower frequency of resistance in that dataset. Of these, 27 were also identified in the RefSeq analysis (Table S17). Most of the other are of unknown function or related to MGEs. Among all the genes of known or putative function that were identified as affecting the emergence of the resistance mutations, we found a few recurrent broad functions involved in mitigation of the effect of quinolones, growth arrest, metal metabolism, biofilm formation, and sugar metabolism.

#### Mitigation of the effect of quinolones

The loss of the *marR* gene is associated with an increase in the probability of subsequent acquisition of resistance in the RefSeq dataset. Inactivating mutations in *marR* were previously found to be associated with decrease of sensitivity to different antibiotics including quinolones (76). *marR* code for a repressor of *marA*, which code for a transcriptional activator of *acrAB* and *tolC*. Increase in the activity of the AcrAB-TolC multidrug efflux pump has been shown to increase resistance to quinolones (77). Inactivating mutations in *marR* are frequent in *E. coli* clinical isolates resistant to fluoroquinolones (78). In viral region 1 we found a gene encoding DinI-like protein. DinI turns off the SOS responses through inhibition of the RecA coprotease activity (Yasuda et al., 1998). *E. coli* mutants deficient in SOS induction were previously shown to survive longer in the presence of several quinolones, suggesting that induction of the SOS response by quinolones is harmful to wild-type cells (79). Finally, the cluster named sit encodes CrcB, a plasmid-encoded protein whose overexpression increases the supercoiling level of plasmids but also reduce the sensitivity of gyrase and topoisomerase IV temperature-sensitive *E. coli* mutants to nalidixic acid (1^st^ quinolone) (80).

#### Cell growth inhibition

Functions associated with bacterial growth arrest are amongst the most represented functions in the 60 groups of genes. Slow growth decreases the bactericidal efficacy of antibiotics, whether because it increases tolerance or persistence (81). Of note, evolution of tolerance was found to systematically precede the experimental evolution of resistance to ampicillin (82). We found eight clusters encoding toxin-antitoxin (TA) systems (NinE, rac, plasmidic group 1, plasmidic group 2, ydfR, TA1, TA2 and TA3) whose acquisition was inferred to increase the acquisition rate of resistance mutations. TA have been describe to induce persister phenotypes that are highly tolerant to antibiotics, including quinolones (83), although recent data has questioned this (84). Three of the identified TA hits are from the ParDE family, which was described to contribute to more than a 1000 fold increase in survival in the presence of supra-MIC concentrations of quinolones (85). Four groups encode cell division inhibitors (*dicC*, *dicB*, *kill* and *rac*). Single-cell imaging showed that ofloxacin persisters formed polynucleoid filamentous cells. This phenotype was independent of the conserved filamentation inducer genes *sulA* or *damX*, suggesting that it was controlled by other cell division inhibitors (86). Interestingly, it was previously found that cryptic prophages of *E. coli* contribute significantly to resistance to sub-lethal concentrations of quinolone through proteins that inhibit cell division, notably KilR of Rac and DicB of Qin (87), both of which were identified in our analysis. Hence, the presence of TA and cell division inhibitors seems to provide a favourable ground to the acquisition of resistance mutations, presumably because they provide more time for the mutations to emerge under antibiotic therapy.

#### Metal uptake

Five clusters encode genes encoding proteins involved in the uptake of metals, presumably iron, zinc, manganese, or copper. In all cases, the acquisition of these genes is associated with a subsequent increase in the rate of acquisition of quinolone resistance. Notably, four clusters (fec, sit, silver, irp) encode almost exclusively genes involved in metals intake, showing that the effect is not related with genetic linkage to other functions. In viral region 2 there is a gene encoding a protein with a domain TonB/TolA, C-terminal. This domain is also involved in iron transport (88). The case of iron uptake could be regarded as surprising, because iron is associated with the creation of reactive oxygen species (ROS) that were suggested to add to the toxicity of antibiotics. Yet, iron acquisition was shown to increase the MIC of quinolones in *E. coli* (89) and to promote the acquisition of quinolone resistance (90). Moreover, several projects are considering therapies that combine ciprofloxacin and iron chelators that reduce the iron availability to bacteria to suppress the growth rate of drug-resistant subpopulations (91). Of note, one of the two groups of genes detected only in the AUS dataset consists of yet another iron siderophore systems of the aerobactin pathway of iron uptake (*iucABD* and *iutA*, Table S16) (92). Other metals, especially copper and zinc, were associated with elevated resistance rates against several antibiotics (93). For example, minimal selective concentration for ciprofloxacin resistance increased up to five-fold in the presence of zinc (94).

#### Biofilm formation

We found three clusters including genes involved in biofilm formation. Of these, the acquisition of two clusters, ag43 and plasmidic group 1, is associated with increase probability of acquisition of resistance mutations. These clusters contain the *Ag43* gene and the *epsB*/*macA*/*macB* genes, respectively, which enhance biofilm formation (95–98). The third cluster contains only the chromosomal gene *yliE/pdeI*, whose over expression reduces biofilm formation (99) and whose mutant formed more biofilm in the mouse gut (100). In our analysis, the loss of this cluster increases the acquisition rate of the quinolone resistance across the tree. Taken together, this suggests that a higher capacity to form biofilms increases the acquisition rate of quinolone resistance. Quinolones are known to be efficient at diffusing through biofilms. (101). However, when compared to planktonic lifestyle, bacteria in biofilms developed more mutants with low-level resistance to quinolones. This is because the biofilm growth mode promotes the upregulation of efflux pumps (102). Even if these low-level resistances do not reach the clinical resistance level, biofilms might increase the time of survival of bacteria, giving them more opportunities for the emergence of mutations responsible for high-resistance levels to quinolone (103, 104).

#### Sugar metabolism

Three clusters contain genes involved in sugar metabolism. Of these, two clusters, *sgc* and *gatC*, contain genes involved in the galactitol metabolism. The gain of *sgc* increases the rate of acquisition of the quinolone resistance, while *gatC* reduces it when lost. A recent screen showed that mutants in the galactitol pathway repressor have a reduced susceptibly to fluoroquinolone in *Salmonella*, suggesting that this pathway is associated with low-level resistance to the antibiotic (105). The third cluster contains the *rfbB* gene that is involved in the dTDP-rhamnose biosynthesis (106). Intracellular dTDP-rhamnose concentration correlates with the MIC of nalidixic acid and norfloxacin (107). This occurs because dTDP-rhamnose upregulates *gyrA* transcription in *E. coli* in the presence of Norfloxacin and nalidixic acid. This helps cells to cope with the quinolones by sequestering the antibiotic and thereby reducing its effect. Taken together, these results suggest that the metabolism of certain sugars may facilitate the acquisition of quinolone resistance by contributing to reduced susceptibility.

### Co-occurrence of the quinolone resistance with other antibiotic resistance classes

We have shown above that *E. coli* strains encode numerous ARGs (Figure 2A, Table S5). Among the 490 RefSeq genomes with point mutations conferring resistance to quinolone, 400 also encode well-known determinants of resistance to beta-lactams, 372 to aminoglycosides, and 352 to sulfonamides (Figure5, Figure S11). This raises the question of how the presence of one antibiotic resistance influences the acquisition of another. We screened our chronologies for antibiotic resistance genes systematically acquired before the resistance mutations and found none that was below the statistical significance level. Hence, it seems that in lineages resistant to quinolones, the acquisition of resistance to quinolones occurs early on, often being among the first resistance mechanisms acquired.

We then checked if other classes of ARGs were systematically acquired after the acquisition of the quinolone resistance mutations. We found 43 ARGs they were significantly more likely to be acquired after the emergence of quinolone resistance (Figure 5, Table S18, Figure S12). For these families, the acquisition of resistance to quinolone increased the rate of subsequent acquisition of the ARG (λ>1). These families correspond to various antibiotic resistance classes among which the most diverse are for resistance to aminoglycosides (12/43), phenicol (6/43), beta-lactamases (5/43), and trimethoprim (5/43). Hence, resistance to quinolones seems to precede resistance to most other antibiotics. Alternatively, quinolone resistance may be less frequently lost and thus remain for longer periods in lineages than resistances that are very costly and/or encoded in MGEs that are easily lost.

**Figure 5:**
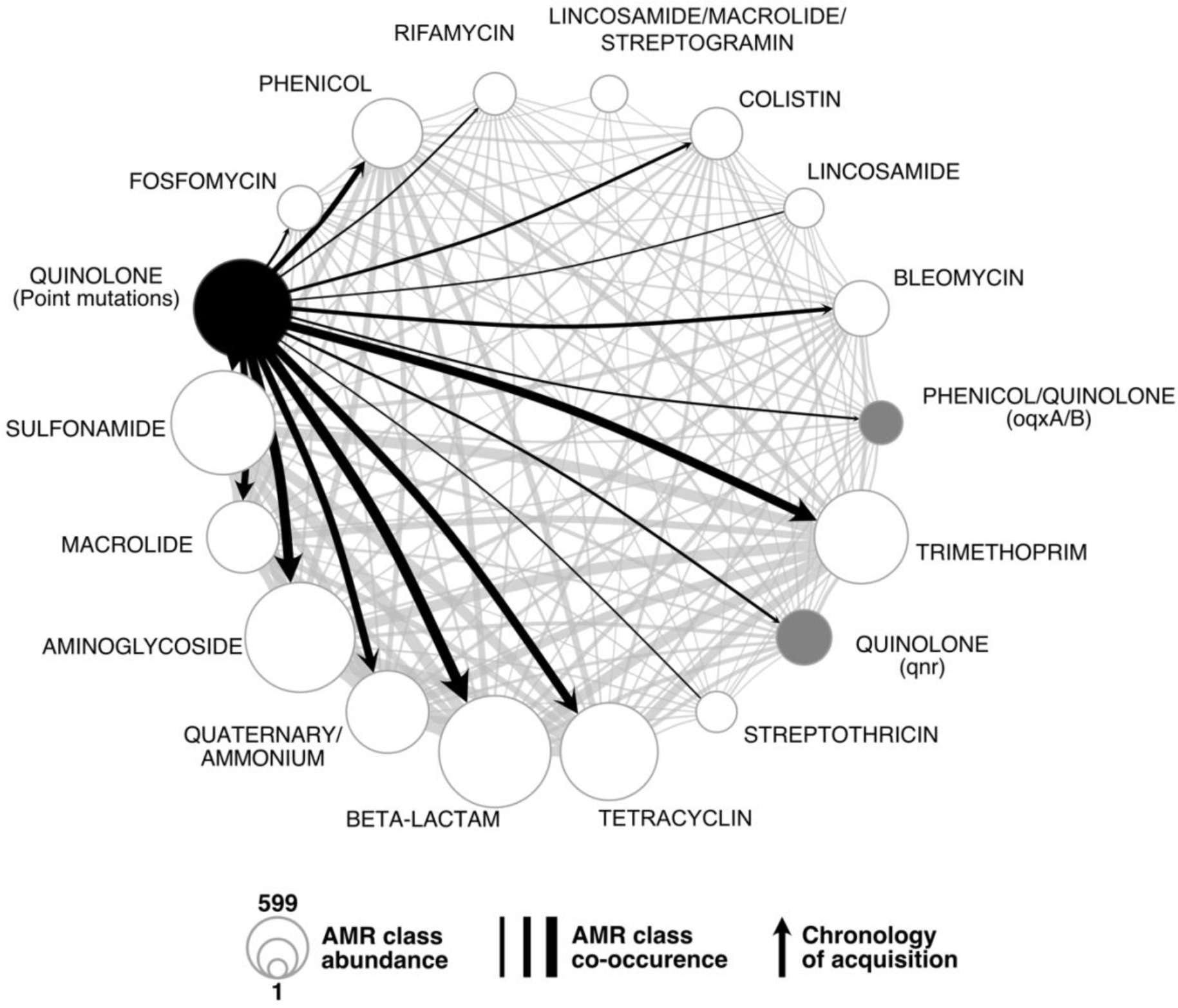
Co-occurrence of the different AMR classes (as defined by AMRfinder+) and the point mutations conferring the resistance to quinolone in the RefSeq dataset. The node size represents the number of times the resistance was found across the genomes. The resistance to quinolone conferred by point mutations and by *qnr* or *oqx* genes are represented in black and grey respectively. The edge width represents the number of co-occurrences in a genome. Edges were converted into arrows when the chronologies between acquisition of mutations and the AMR classes were significant (mutations emerging first). When the resistance is conferred by other mechanisms, genes involved are annotated between brackets.

As mentioned above, low-level resistance to quinolones can also be acquired through the acquisition of certain plasmid-associated genes. Interestingly, accumulation of these resistance mechanisms is frequent. Three of the families identified as systematically being acquired after the point mutations conferring resistance to quinolone correspond to genes that provide low-level resistance to quinolones (either *qnr* or *oqx*). Hence, we observe an initial emergence of point mutations conferring fluoroquinolone resistance and then the acquisition of specific resistance genes by horizontal gene transfer. This is in line with historical data: resistance to quinolones due to point mutations is almost as old as the quinolone usage. In contrast, resistance due to acquired genes is more recent. This accumulation is clinically relevant because the resistance effect provided by these novel genes affect resistance to all quinolones (at least for *qnr*) and it is additive to the one provided by the point mutation (resulting in increased resistance) (108).

## Discussion

In this study we have searched to identify chronologies shedding light on the evolution of resistance to quinolones in *E. coli*. We have searched for them before, during, and after the acquisition of resistance. We found many genes that were preferentially gained/lost before the acquisition of the resistance mutations. During the process of acquisition of the mutations there is a preferred initial one and then a very quick succession of mutations whose order we could not trace. After the acquisition of resistance, we found that other resistances were often acquired after quinolone resistance mutations. It should be emphasized that these chronologies correspond to pairs of events that are far apart in time (they are found in different branches of the phylogenetic tree). They should not be interpreted in the framework of hysteresis, which is a transgenerational change in cellular physiology (109). Instead, we interpret the chronologies for gene gains/losses as changes that pre-dispose a lineage to the acquisition of resistance, but that were selected for other reasons. We think many of the genes identified by this approach fall into three categories: functions in epistatic interaction with quinolone resistance, functions co-selected in lineages for ecological/epidemiological reasons, and genes in genetic linkage with the former.

Many of the functions frequently acquired before the resistance mutations might make the bacterium more tolerant, resistant, or able to persist in the presence of quinolones. This is the case of functions that are known to arrest growth (TAs), those that diminish the availability of quinolones as a side-effect (transporters, metal chelators), or those that increase the expression of the gyrase. These functions could allow the bacterium to cope with the presence of quinolones for a longer period and thus provide time for resistance mutations to arise. This does not require any kind of foresight for natural selection. These functions might have their own adaptive role and may have been fixed independently of the possible advantage they may subsequently give under quinolone treatment. In some cases, the phenotype under selection before antibiotics is also adaptive under antibiotic therapy. In other cases, the phenotypes under selection with or without antibiotics may be very different, as observed for the acquisition of low dose resistance to quinolones by over-production of gyrase by the dTDP rhamnose (107). Such latent phenotypes may be frequent across microbial systems, as previously suggested for gene regulatory circuits (110), moonlighting proteins (111), and biofilm formation (112). The constant gain and loss of genes in bacterial genomes may produce a wide diversity of such latent phenotypes in natural populations rendering certain lineages better prepared than others to acquire specific novel traits (like antibiotic resistance). Such pre-disposition does not preclude the acquisition of resistance in other lineages that did not acquired these genes, because the trait is accessible by mutation and is under strong selection. What we propose is that acquisition of certain genes might increase the probability of subsequent emergence of resistance to quinolones either because they increase the likelihood of mutations or because they provide less costly genetic backgrounds.

Epistatic interactions in the process of accumulation of the mutations conferring resistance to quinolones have been described before (29, 113). The chronology usually starts by the mutation *gyrA*[S83L] (which was shown to have the largest effect and low cost) followed by quick fixation of the *parC*[S80I] and *gyrA*[D87N] mutations, which further increase resistance at low cost (20, 29, 55, 70, 114). The second most frequent evolutionary trajectory is the simultaneous acquisition of the three mutations. We thought this could be due to recombination, since the simultaneous acquisition of 4 mutations in *gyrA*, *gyrB*, *parC* et *parE* after one single event of chromosomal conjugation was described in mycoplasma (115). Yet, we detected only one case with clear evidence that the mutations in *gyrA* and *parC* were acquired in the same branch of the tree by recombination. Alternatively, the lack of time-resolution at the scale of the species trees could explain the apparent simultaneous emergence of all mutations. To test this idea, we performed the same analysis on three STs for which many genomes were available, which should improve the resolution of our study. Even in this case, we failed to untangle the order of acquisition of the mutations. While this matter remains to be completely resolved, we propose that natural selection for resistance by multiple mutations may be so strong in the clinical environment (14, 33) that the multiple point mutations are quickly acquired in succession and outcompete the other mutants. If so, then these lineages have frequently endured three successive (possibly nested) selective sweeps.

Epidemiological factors may explain some of the identified chronologies. The lineages of pathogens are more likely to encounter antibiotic pressure. Mutations conferring the resistance to quinolone are then more likely to be selected in such lineages. Hence, the acquisition of virulence factors may be associated with an increase in the rate of emergence of quinolone resistance once it started to be used in the clinic. Of note, the most obvious virulence factors (such as toxins, protein secretion systems or their effectors) were not revealed by our analysis. But we did find functions that while being very frequent in environmental bacteria are also associated with host colonization, like biofilm formation or iron uptake. Although there is data associating these traits with the acquisition of quinolone resistance outside the context of pathogenesis (90, 116), one cannot exclude the possibility that some of our results are due to epidemiological factors.

Epidemiological factors seem particularly pertinent to explain the chronologies of ARG acquisitions after the emergence of quinolone resistance. None of the genes systematically acquired before the quinolone resistance was annotated as ARG. However, genes encoding resistance to many antibiotic classes such as beta-lactams and aminoglycosides are consistently acquired after quinolone resistance. This suggest that the quinolone resistance was usually acquired before they evolved to become multi-drug resistant. Historically, resistance to quinolone was not the first one acquired by *E. coli* where resistance to penicillin was described in the 40’s while quinolone resistance only in the 70’s (117, 118). However, quinolone resistance is the one that best correlates with its antibiotic usage in *E. coli* across European countries (42), in agreement with this and other works showing that it can emerge very rapidly (78). The accumulation of the full set of resistance mutations makes the clones fitter when compared to other antibiotic resistances so that the resistance is less likely to be lost (20, 70). It may also be difficult to lose the resistance to quinolones by multiple mutations since it requires the reversion of all of them, whereas many other resistances may be lost simply by plasmid loss, gene deletion or inactivation. Several studies have reported that mutations in the core genome often do not reverse even when antibiotic resistance pressure was removed (72, 119, 120). This combination of high acquisition/low reversion rates could be the reason why the quinolone resistance appears very early in lineages of bacteria becoming multi-drug resistant. Nevertheless, some studies have shown that resistance to quinolones does promote resistance to other antibiotics. For example, substitution at *gyrA*[87] in Salmonella influences sensitivity to other types of antibiotics and results in increased expression of stress response pathways (121). Hence, even the chronologies of gene gains/loss after acquisition of resistance may result from epistatic interactions.

A specificity of our approach is that functions acquired as the result of the same event of transfer will have similar statistical signal. Yet, this does not mean they are all biologically pertinent. One single gene under strong selection and strong linkage with others may lead to the entire set of genes coming out as significant in our analysis. Our clustering approach allowed to regroup such genes together and provides a more parsimonious analysis of the functions that may explain the chronologies. Several of these groups only have genes of unknown function. It’s beyond the scope of this work to elucidate these functions or to disentangle which is most important in each group of linkage. Yet, our analysis reveals many functions that were previously shown to be involved in the acquisition of quinolone resistance (or associated with it). This validates our approach and provides a rich list of genes for future experimental analysis.

The clustering of genes in linkage also allowed to study the vehicles of gene acquisition, i.e. the mobile genetic elements that promoted the gene transfer. Several of these groups of genes are in plasmids or prophages. If the acquisition of one gene carried by an MGE influences the acquisition of the mutation conferring resistance due to epistatic interactions, other genes carried by the same MGE will hitchhike with it. This has often been theorized as being one of the potential reasons why costly and autonomous MGEs, such as prophages and conjugative elements, encode traits adaptive for bacteria (122). Hence, we do not think that the entire MGEs are favoring the acquisition of quinolones, only a few of their genes. Mobile genetic elements, especially plasmids, are known to carry ARGs to the bacteria they infect. But our results suggest they are also responsible for facilitating the acquisition of antibiotic resistance by carrying functions that make the bacterium better prepared to acquire the resistance itself. In this context, the identification of prophages having such an effect is intriguing (even though it fits previous works (87)). Quinolones are known to induce the SOS response which induces the lytic cycle of many prophages (123). It’s tempting to speculate that prophages might carry traits favoring resistance or tolerance to quinolones to avoid induction at a moment where most potential hosts in the near environment are not promising hosts for infection. These genes may not have evolved to tackle quinolone resistance, presumably a recent threat to *E. coli*, but may have become advantageous under it. Anti-SOS genes have been found in both conjugative plasmids and prophages where they may serve to manipulate the host responses or those of other mobile elements (124, 125). This fits our observation that the prophages identified in this work encode functions resulting in growth arrest or mitigation of the effect of quinolones.

Bacteria colonize diverse environments that vary in terms of selective pressures. Epistatic interactions create complex evolutionary patterns of adaptation. Our results suggest that this complexity further increases when a mix of horizontal gene transfer and mutations produce bacteria with very different genetic backgrounds within lineages. While these interactions can facilitate the emergence of antibiotic resistance, they may also render evolutionary processes less predictable (126). They may contribute to explain why within many microbial species some lineages tend to accumulate traits like virulence factors and antibiotic resistance. Interestingly, recent studies show that the decline of resistance to a cephalosporin in *Pseudomonas* is also contingent on the genetic background (127). Understanding how frequent chronologies of events result from genetic, functional, and epidemiological interactions will shed further light on the evolution of antibiotic resistance and how one can forecast it.

## Methods

### Genome data and pangenome construction

We analyzed three datasets in this study.

The first dataset is the one used across the study unless otherwise specified. It includes 1585 genomes of *E. coli* identified from 21 086 complete bacterial genomes retrieved from NCBI RefSeq representing 6 124 species of Bacteria (http://ftp.ncbi.nih.gov/genomes/ <u>refseq/bacteria/</u>), in March 2021. This dataset has the advantage of being very large, diverse, and we know the structure of the genome (chromosome, plasmids). This facilitates the study of the mobile genetic elements allowing gene acquisition, such as plasmids and phages, and provides high quality data for the identification of mutations. It is usually presumed that it contains many clinical isolates.

The second dataset was used to control for the presumed over-representation of clinical isolates in the RefSeq dataset. It includes a collection of 1 294 *E. coli* draft genomes from isolates recovered in Australia between 1993 and 2015 chosen to represent the phylogenetic diversity of the species (69). It includes less than 25% of clinical isolates and allows to control for sampling biases commonly encountered in public databases.

The 2 879 *E. coli* genomes were analyzed for assembly quality using their L90 value and for genetic distance using MASH v2.2 (128). We removed from further analysis 369 strains from the Australian dataset because they had a L90 superior to 100 (i.e. the sum of the 100 longest contigs does not cover at least 90% of the genome size, suggesting that they are highly fragmented). Additionally, 365 were removed because they were very similar (MASH distance <10^-4^). This resulted in a dataset of 2 145 (877 from the Australian dataset and 1 268 RefSeq sequences) completely sequenced and assembled *E. coli* genomes.

The third dataset was only used to pinpoint the chronology of mutations for quinolone resistance at a short evolutionary time scale. We used all *E. coli* sequence type 131 (ST131) genomes from RefSeq (n=85) and we retrieved all available *E. coli* sequence type 410 and 167 (ST410 and ST167) genomes from Enterobase (n=1000 and n=801) (last accessed in December 2021)(129). We removed from further analysis 45 ST410 and 29 ST167 genomes because they had a L90 superior to 100. Additionally, 198 ST410 and 193 ST167 strains were removed because they were too similar (MASH distance <0.0001). This resulted in a dataset of 757 *E. coli* ST410, 85 *E. coli* ST131 and 579 *E. coli* ST167 genomes.

### Identification of the pangenome and of the core genome

The pangenome and core-genome of the 2 145 *E. coli* from the RefSeq and Australian dataset were computed using PanACoTA v.1.3.1 (130). Briefly, the pangenome was constructed by clustering all protein sequences in the set of genomes using MMseqs2 (protein identity >80%). We retrieved the core-genome from this pangenome. It consists of the 2393 gene families present in exactly 1 copy in at least 99.0% of all genomes. In the remaining 1% genomes, we accepted the presence of 0 or several members of the gene family (in which case the sequence of the gene is not used for the strain). The pangenome (27556 gene families) and core-genome (3933 gene families) for the 757 ST410 *E. coli* were built the same way.

### Phylogenetic reconstruction

The multiple sequence alignments of the families of the core genes were computed using the ‘align’ PanACoTA v1.3.1 module. Briefly, the protein sequences of the core genes were aligned with MAFFT v7.467 (-auto parameters) (131) then back-translated to nucleotide alignments (*i.e.*, each amino acid was replaced by the original codon) and concatenated. The large multiple sequence alignment including the RefSeq and Australian dataset was then separated in an alignment with the 1 268 RefSeq genomes and another with the 877 genomes from the Australian dataset. The phylogenetic inference was done from the resulting multiple alignments using IQ-TREE 2.0.6 (132) with the ultra-fast bootstrap option (-bb 1000 bootstraps) and with the best fitting model estimated using *ModelFinder Plus* (-MPF) (133). The best model for the three datasets was GTR+F+I+G4 according to the BIC criterion. Trees were rooted using the midpoint function from the phangorn packages (v.2.5.5) for R (134). The root obtained is in agreement with the literature, as strains belonging to Clade 1 (outgroup closest to the E. coli species) are indeed the most external.

### Mutation profile

Point mutations leading to quinolone resistance in *E. coli* are found in the DNA gyrase and the DNA topoisomerase IV genes, i.e., in *gyrA, gyrB*, *parC* and *parE* (45). We retrieved the sequences of these proteins using blastp v2.12.0 (default parameters, identity threshold of 90%) (135). We then built a multiple alignment of the proteins using MAFFT v7.429 (with - auto parameters). The alignments were parsed using Biopython (136). We looked for point mutations leading to quinolone resistance, at each expected position (accounting for gaps) using the comprehensive list provided by Hopkins et al. (45) for *E. coli* (Table S1). For each dataset, the distribution of every combination of resistance mutation was summarized in an upset plot computed using the R package UpsetR v1.4.0.(137)

### Inference of ancestral gene repertoires

We counted the number of occurrences of each family of the pan-genome in all genomes. This was used to build a gene presence (1 or more copies)/absence matrix in all leaves of the phylogenetic trees. From this occurrence matrix, we inferred the ancestral state (presence or absence) of each gene family at every internal node of the phylogenetic trees with PastML v1.9.33 (138). We used the JOINT method with default parameters, this method reconstructs the states of the scenario with the highest likelihood. From the ancestral state matrix computed by PastML, we inferred the gene gains and losses at all the branches of the species tree by subtracting the gene content of the child node to the gene content of the parent node.

### Trajectories of acquisition of mutations

To infer the history of the quinolone resistance mutations, we constructed a matrix with the presence or absence of each type of mutation in *gyrA*, *gyrB*, *parC* and *parE* in each strain. The ancestral states for each mutation were inferred the same way as described in the previous section for the genes in the pangenome. To identify the acquisition chronologies of the multiple mutations, every path starting from the root and leading to a leaf containing at least one quinolone resistance mutation was extracted using the ape package v5.3 for R (139). Paths were then traversed from the root to the first acquisition of a mutation conferring quinolone resistance (as inferred by PastML). All paths arising from this first mutation were then traversed from this event of acquisition to the next event on the path and this recursively until the last event of acquisition on each path (for the scripts used to perform this analysis, see supplementary files). This way, we only consider the number of events in the tree, not the number of taxa affected by them. This is important because events of gain and loss within a gene family can be regarded as independent, whereas resistant taxa are not (they may result from the same ancestral mutation event). Counting resistant taxa would inflate and bias the statistics, over-representing some chronologies. Counting events allows to identify independent events across the tree. This procedure results in a collection of paths corresponding to every single chronology of mutation acquisition.

### Detection of preferential chronologies

Preferential chronologies of events leading to the acquisition of resistance to quinolones were identified in a two-step procedure using the program Evo-Scope v1.0.0 (140).

In the first step, we analyze all possible chronologies to identify events often preceding the acquisition of resistance to quinolones. This is done with the Epics module of Evo-Scope (with the -S parameter). The procedure identifies the number of occurrences of an event E1 (e.g., acquisition of resistance) following an event E0 (e.g., acquisition of a certain gene) on a phylogenetic tree. The observed values of all pairs of events are compared with the expected numbers under a null model of uniform rates of distribution of events on the tree (75). Pvalues were adjusted for multiple tests using the ‘fdr_bh’ method (P<10^-5^) (Benjamini-Hochberg correction) from SciPy v.1.10.1 (141, 142).

In the second step, we retrieved the significant chronologies identified above and inferred the interaction type and strength of correlated evolution between these pairs using the Epocs module of Evo-Scope (143). This module classifies interactions, by maximum likelihood, in 3 different categories depending on the influence of a trait on the occurrence of the other one.

**- Scenario of independence (-Si):** The occurrence of event E0 does not change the occurrence rate of E1.
**- Scenario of asymmetric induction (-Sa, -Sb):** The occurrence of event E0 changes the occurrence rate of event E1 or the occurrence of event E1 changes the occurrence rate of event E0.
**- Scenario of reciprocal induction (-Sl):** Event E0 enhances the occurrence rate of event E1 and, reciprocally, event E1 enhances the occurrence rate of event E0.

These scenarios are described by models containing from two to four parameters, which are associated to each trait. The parameters are divided into natural occurrence rates (i.e., rates at which the trait mutates from present to absent or from absent to present) and modified occurrence rates (i.e., rates at which the trait mutates from present to absent or from absent to present after a change of state of the other trait). The ratio λ between the modified occurrence rates and the natural occurrence rates can be interpreted as an induction factor. If λ > 1, the induction between the two traits is positive (i.e., E0 increases the occurrence rate of the subsequent event E1) whereas it is negative when λ < 1 (i.e., E0 decreases the occurrence rate of the subsequent event E1). The model best describing the data under study is selected following a significant Likelihood Ratio Test (144)

### Detection of recombination

The alignment of the core-genome that was used to build the phylogeny was scanned with Gubbins v2.4.1 to find recombination tracts (145). The maximum likelihood tree previously built was provided for the first iteration using the “—starting-tree” command. The position of the recombination tracks was compared with the position of mutations in *gyrA* (*gyrA*[S83L] and *gyrA*[D87N]) and in *parC* (*parC*[S80I]) in the alignment of the core-genome to identify overlaps that could indicate that mutations arrived by recombination. We then compared the nodes at which recombination occurred with nodes where the mutations *gyrA*[S83L], *gyrA*[D87N] and *parC*[S80I] mutations were acquired. When overlapping recombination tracks and acquisition of mutations occurred at the same nodes, we considered the mutations to be acquired by events of recombination.

### Genome-wide association study

We performed a genome-wide association study (GWAS) for the presence of fluoroquinolone resistance using 1268 *E. coli* genomes with pyseer v.1.3.9 (64). The association between the gene presence/absence and the resistance phenotype (defined by the presence of a known resistance mutation) was assessed with a linear mixed model (LMM) in which the Sequence Types (ST) were considered as covariates. We used the recombination-free phylogenetic tree produced by Gubbins. This tree allowed us to generate a distance and a kinship matrix with the scripts coming with pyseer. The linear-mixed model used the multidimensional scaling (MDS) of these matrices to control for population structure. 10 dimensions were included in the MDS. To address the problem of multiple comparisons in our analysis, we used a Benjamini-Hochberg procedure (141) on the p-value of the association already adjusted for population structure using the p.adjust function with the “BH” method in R v4.2.2. For a corrected p-value inferior to 0.05 we deemed the association between the gene presence/absence and the resistance as significant.

### Identification of groups of genes in genetic linkage

Gene families consistently acquired before the resistance to quinolone can be in genetic linkage, *e.g.,* if they are systematically co-acquired within a plasmid, a phage or recombination. In such cases, there may be only one gene that effectively changes the likelihood of acquisition of the resistance, but the method will also highlight genes in strong linkage with this one. To control for this effect, we took all the genes highlighted by the analysis of chronologies and clustered them using two key information: first co-acquisition or co-loss in the phylogenetic tree and then co-localisation in the genome.

Firstly, two gene families were clustered together if they were consistently co-acquired or co-lost in the same branch or node of the tree. To assess statistical significance of these simultaneous events, we used the Epics module of the program Evo-Scope with the parameter -I (i.e, Identity matrix). Thus, the program compares the number of co-occurrences of two events E1 and E0 in a branch of the tree to their expected co-occurrence under a null model of uniform distribution of events on the tree. Pairs of events that frequently co-occurred in time were then clustered by single-linkage using the agglomerative clustering algorithm from scikit-learn v1.2.2 (parameters: affinity=’precomputed’, distance_threshold=1, linkage=’single’, n_clusters= None) (146). We used single-linkage to obtain large clusters that could then be further split.

Secondly, we split the clusters of co-acquired or co-lost genes using information on the distance between the genes in the genomes. For every pair of gene families in a cluster of co-occurrences, we computed the median number of genes between them in the genomes (in which they co-occur). As all pairs of genes need a distance, when genes were present on different DNA molecules (e.g., one in the chromosome and another in a plasmid), the number of genes in the largest replicon in both bacteria was set as the distance between them. This way, the distance between genes on different DNA molecules will be above the clustering threshold (they will not be clustered together). These values were then used to cluster the gene families by average-linkage using the agglomerative clustering algorithm from scikit-learn v1.2.2 (parameters: affinity=’precomputed’, distance_threshold=30, linkage=’average’, n_clusters=None). At the end of these procedures all clusters were checked for homogeneity of induction values (λ) and event type (loss or gain).

### Functional annotation of the pangenome families

We picked at random a representative sequence of each pangenome family. These sequences were annotated using eggNOG-mapper v2.1.9. In order to be exhaustive, we also fetched the gene name and product functions from the RefSeq annotations. When both eggNOG and RefSeq yield different gene names, we used the RefSeq gene name.

### Detection of the antibiotic resistance pangenome families

We picked at random a representative sequence of each pangenome family. These sequences were screened for antibiotic resistance genes using AMRfinderPlus v3.10.18 with default parameters (147). If the representative sequence was identified as an AMR gene the pangenome family was considered as an antibiotic resistance one.

### Statistics

Unless mentioned otherwise all statistics were performed within R (v3.6.3).

## Acknowledgements

Eugen Pfeifer for providing the viral regions data. Manuel Ares-Arroyo for scientific discussions. This project was funded by the INCEPTION project Path2Resistance [PIA/ANR-16-CONV-0005], Equipe FRM [Equipe FRM/EQU201903007835], Laboratoire d’Excellence IBEID [ANR-10-LABX-62-IBEID], SEQ2DIAG [ANR-20-PAMR-0010]. This work used the computational and storage services (MAESTRO cluster) provided by the IT department at Institut Pasteur, Paris. We thank Frédéric Barras, Jessica El Khoury, and Laurence Van Melderen for help on the interpretation of our results.

## Author Contributions

Design of the study: CC, MGu, PG, EPCR

Performed the analyses: CC, MGu, FM

Contributed with data or methods: MT, MGo, GA

Interpreted the analyses: CC, MGu, EPCR

Wrote the draft: CC, EPCR.

All authors participated during writing of the manuscript and approved the submitted version.

## Notes

### Competing Interest Statement

The authors have declared no competing interest.

