## Supplemental Materials for "Chance favors the prepared genomes: horizontal transfer shapes the emergence of antibiotic resistance mutations in core genes"

**This file includes:**

Figures S1 to S12

Supplementary tables

Scripts used in the study

Tree files in newick format

### Supplementary table legends:

**ST1: List of the mutations known to confer resistance to quinolone used as reference.** Each line corresponds to a mutation, indicating the gene concerned, the original amino acid, the position of the mutation, and the amino acid conferring the mutation.

**ST2: List of the mutations conferring resistance to quinolone per genome in the RefSeq dataset.** Each line corresponds to a genome, indicating its strain name, assembly ID and the detected mutations conferring resistance to quinolone.

**ST3: List of the mutations conferring resistance to quinolone per genome in the Australian dataset.** Each line corresponds to a genome, indicating its strain name and the detected mutations conferring resistance to quinolone.

**ST4: Isolation source of the Australian genomes.** Each line corresponds to a genome, indicating its unique name, source of isolation, genome size and phylogroup.

**ST5: List of the antibiotic resistance classes per genome in the RefSeq dataset.** Each line corresponds to a genome, indicating its strain name, assembly ID and the antibiotic resistance classes it encodes as defined by AMRFinder+.

**ST6: List of the trajectories of acquisition of the main resistance mutations in E. coli for the RefSeq dataset:** Each line corresponds to a unique trajectory of acquisition of the mutations of resistance and the number of times it is observed in the tree.

**ST7: List of the trajectories of acquisition of the main resistance mutations in E. coli for the Australian dataset:** Each line corresponds to a unique trajectory of acquisition of the mutations of resistance and the number of times it is observed in the tree.

**ST8: List of the trajectories of acquisition of the main resistance mutations in E. coli for the ST410 dataset:** Each line corresponds to a unique trajectory of acquisition of the mutations of resistance and the number of times it is observed in the tree.

**ST9:** **List of the pangenome categories consistently acquired or lost before the acquisition of the quinolone resistance in the RefSeq dataset.** Each line corresponds to a pangenome category, indicating its name, lambda value, function, Epics and Epocs pvalue, location in the bacteria, EggNog and TAsMania annotations and if it has been gained or lost.

**ST10: List of the pangenome categories found associated with the quinolone resistance by Genome-Wide association study (GWAS):** Each line corresponds to a pangenome category, indicating its pvalue, pvalue corrected for multiple tests, effect on the presence of the mutation, the value of the effect (inferred by pyseer), frequency in the dataset and its EggNog annotations.

**ST11: Matrix used to perform the co-occurrence clustering.** The matrix is an adjacency matrix. Zeros represent co-acquired or co-lost pangenome categories, ones represent those that are not.

**ST12: List of the pangenome categories statically co-occurring.** Each line corresponds to pair of pangenome categories that are statically co-occurring according to epics, indicating the pangenome categories involved in the pair, the Epics pvalue, the number of the co-occurrences observed in the tree and the maximum of possible co-occurrences.

**ST13: List of the pangenome categories consistently acquired or lost before the acquisition of the quinolone resistance in the RefSeq dataset and their cluster of co-occurrences.** Each line corresponds to a pangenome category and its respective cluster of co-occurrences.

**ST14: Median genomic distance between each pair of pangenome categories present at least once in the same genome.** Each line corresponds to a pair of pangenome categories, indicating the number of proteins in each pangenome category, the number of times the two pangenome categories are present in the same genome, and the median distance between them in terms of gene number

**ST15: List of the 60 clusters of genomic co-localization.** Each line corresponds to a cluster of co-localization, indicating its name, lambda mean, effect on the acquisition of the quinolone resistance, main function, location in the bacteria and if it has been gained or lost. The names and functions of all the genes inside the cluster are given in a conserved order.

**ST16:** **List of the pangenome categories consistently acquired or lost before the acquisition of the quinolone resistance in the Australian dataset.** Each line corresponds to a pangenome category, indicating its name, lambda value, function, Epics and Epocs pvalue, EggNog annotations and if it is significantly detected in the RefSeq analysis.

**ST17: List of the pangenome categories consistently acquired or lost before the acquisition of the quinolone resistance in both datasets.** Each line corresponds to a pangenome category, indicating its name, function, lambda value in the RefSeq and in the Australian analysis and the name of the cluster it belongs to in the RefSeq dataset.

**ST18: List of the antibiotic resistance classes consistently acquired after the acquisition of the quinolone resistance.** Each line corresponds to a pangenome category, indicating its name, lambda value, Epics and Epocs pvalue, if it has been gained or lost and its AMRFinder+ annotations

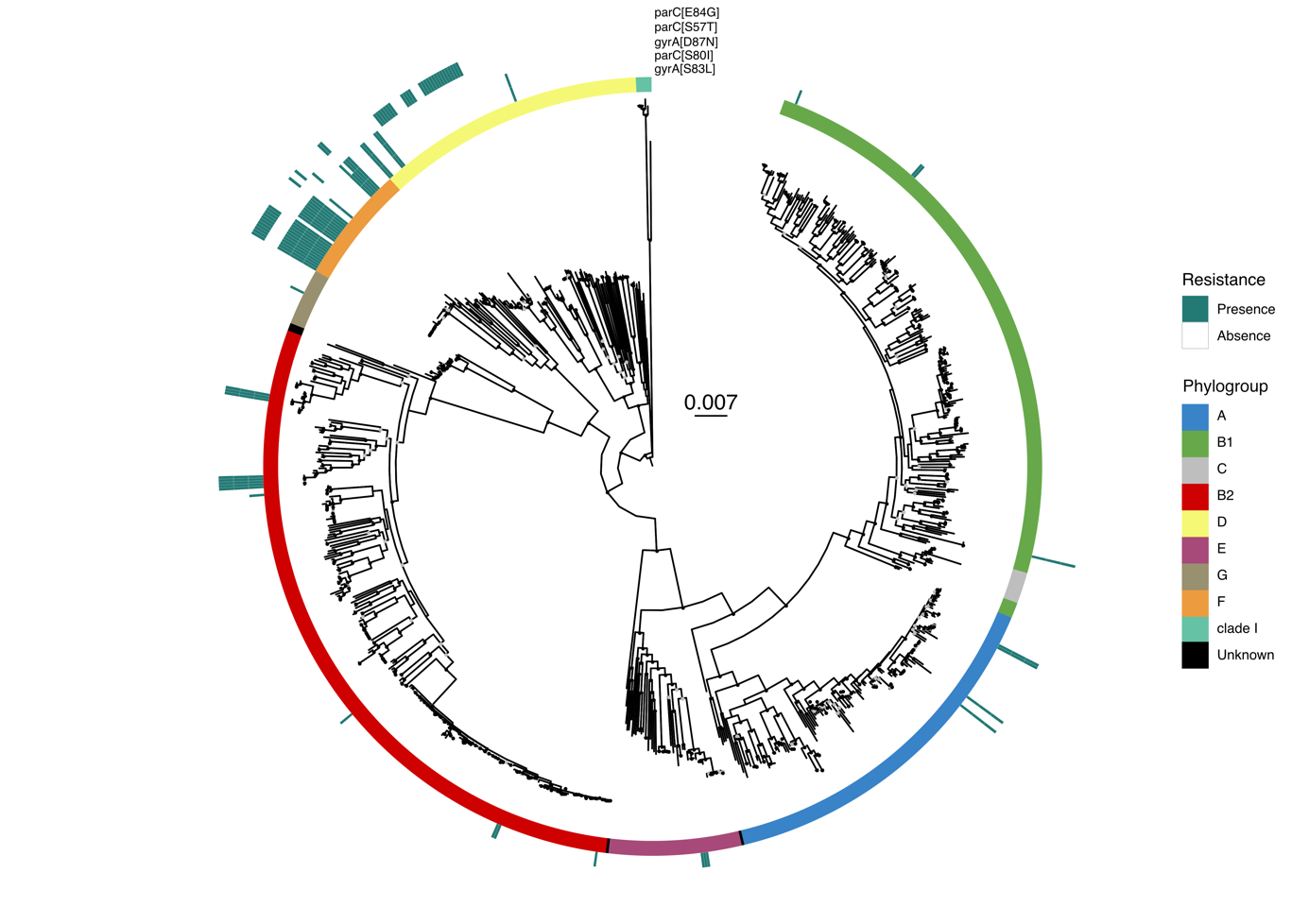

**Figure S1: Phylogeny and distribution of quinolone resistance mutations in DNA gyrase and topoisomerase IV genes for E. coli of the Australian dataset.** The maximum likelihood phylogenetic tree of the species and the scale of substitution per site are in the center with IQTree (model GTR+F+I+G4 and 1000 ultrafast bootstraps). The E. coli phylogroups are represented by the colors in the inner center. The marine green squares indicate the presence of the respective mutation across the species. Ultrafast bootstrap values superior to 75% are shown with a light gray circle and values superior to 90% with a black circle.

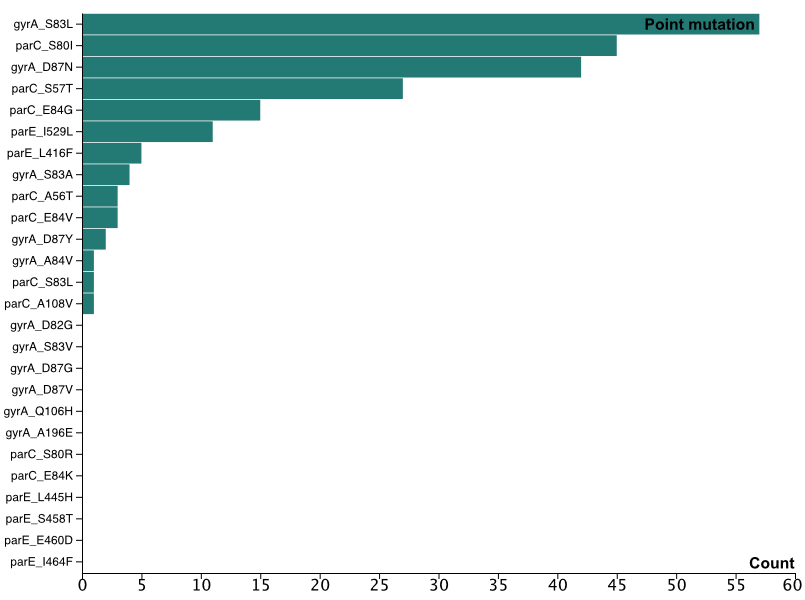

**Figure S2: Distribution of the different quinolone conferring mutations in the Australian dataset.** Mutations are ordered according to their frequency. The first letter in the name after the underscore correspond to the ancestral amino acid, the number to the position of the transition and the last letter to the amino acid conferring the resistance, e.g., gyrA_S83L means that the mutation is in gyrA at position 83 and involves a substitution Ser->Leu.

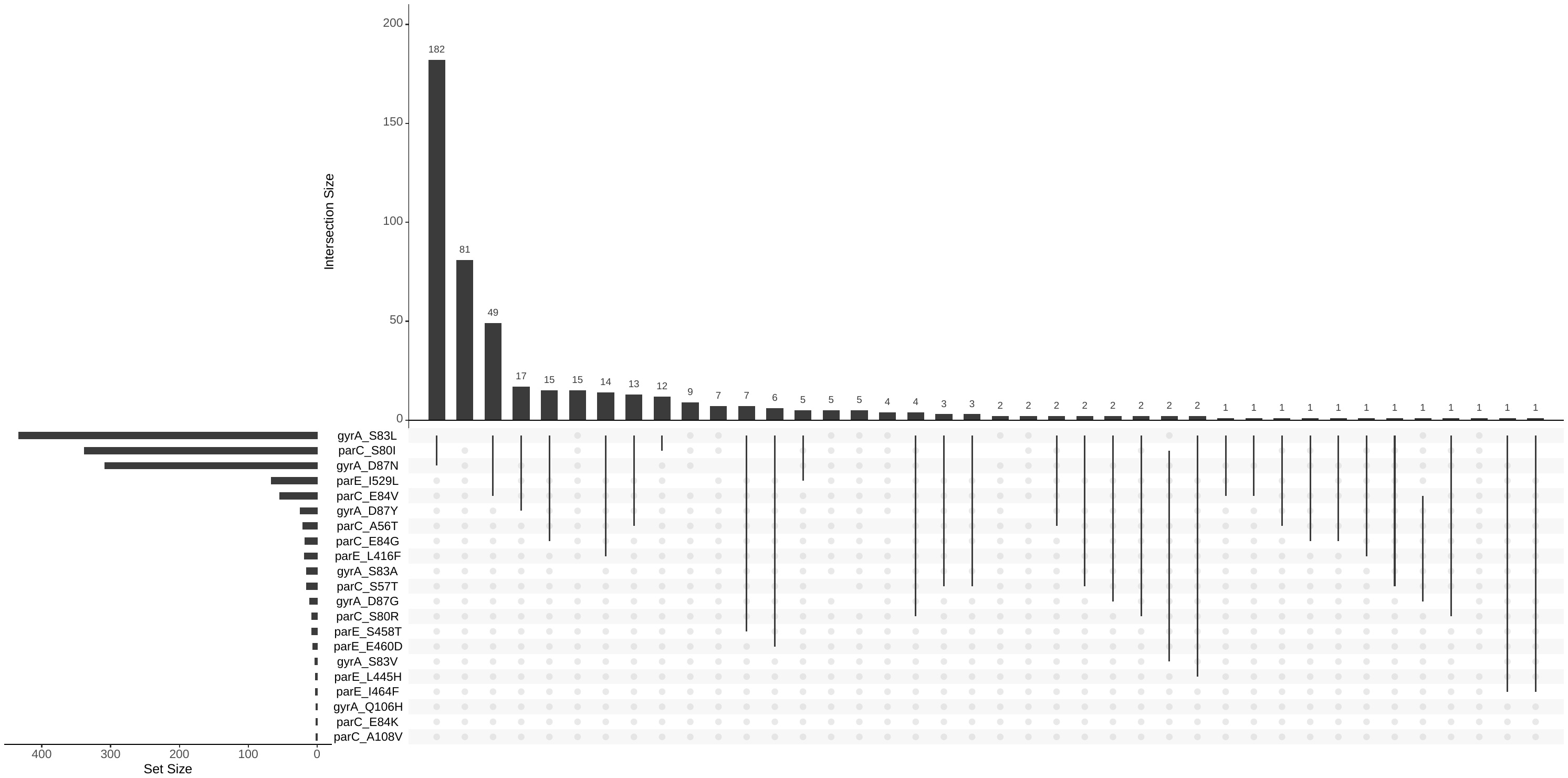

**Figure S3: Upset plot of the different quinolone conferring mutations in the RefSeq dataset.** The horizontal bar plot represents the distribution of the mutations in the RefSeq dataset. The vertical bar plot represents the distribution of the mutation combinations. Lines and dots represent the different combinations.

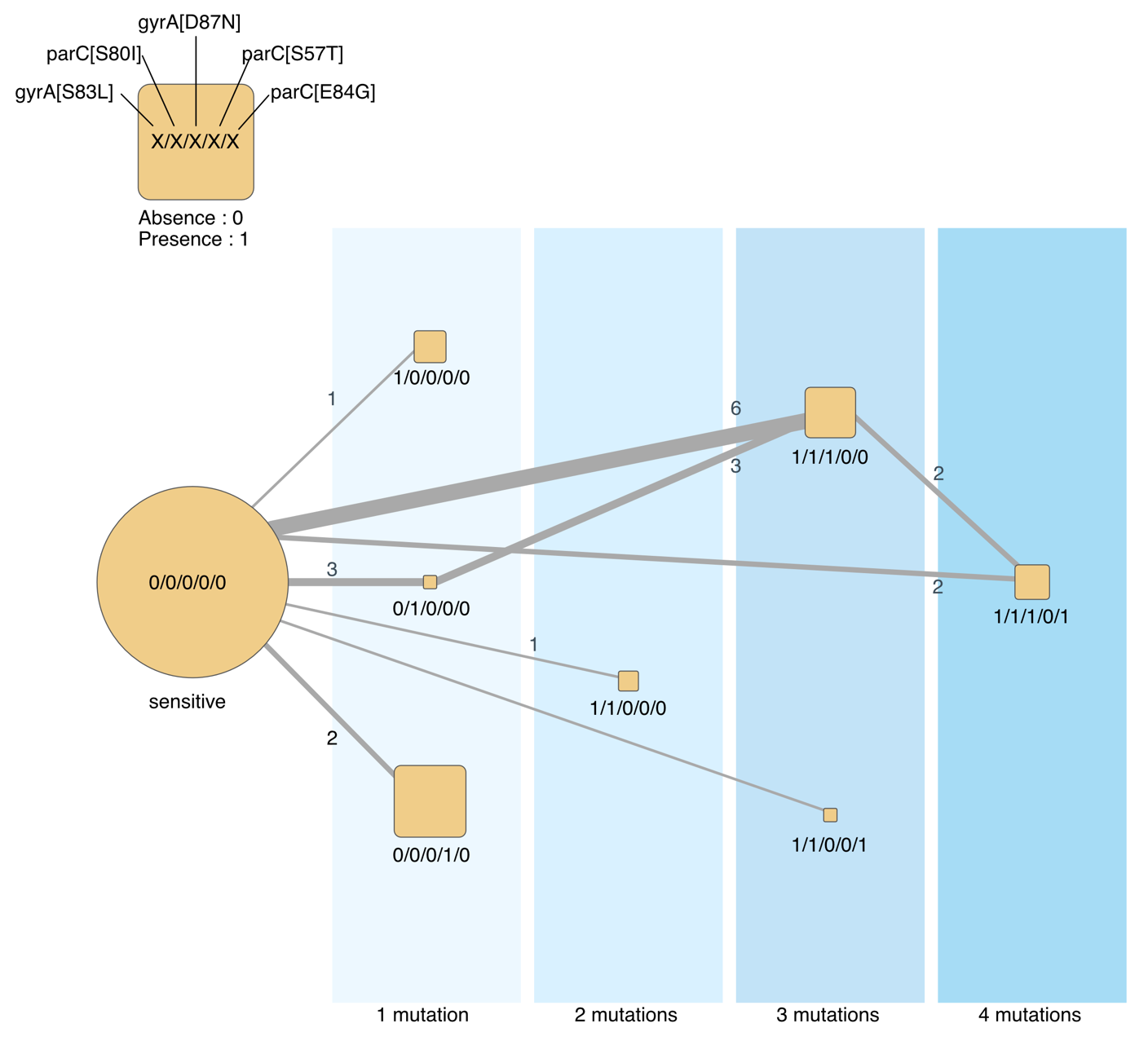
**Figure S4: Trajectories of acquisition of the main resistance mutations in E. coli for the Australian dataset.** The blue areas represent the number of distinct mutations in genomes. The size of the square scales with the number of genomes observed (leaves of the tree). The edges represent the chronologies of acquisition of one or several mutations as inferred from the reconstruction of ancestral states on the tree. The edge size is proportional to the frequency of the respective transition, with the labels showing this exact number.

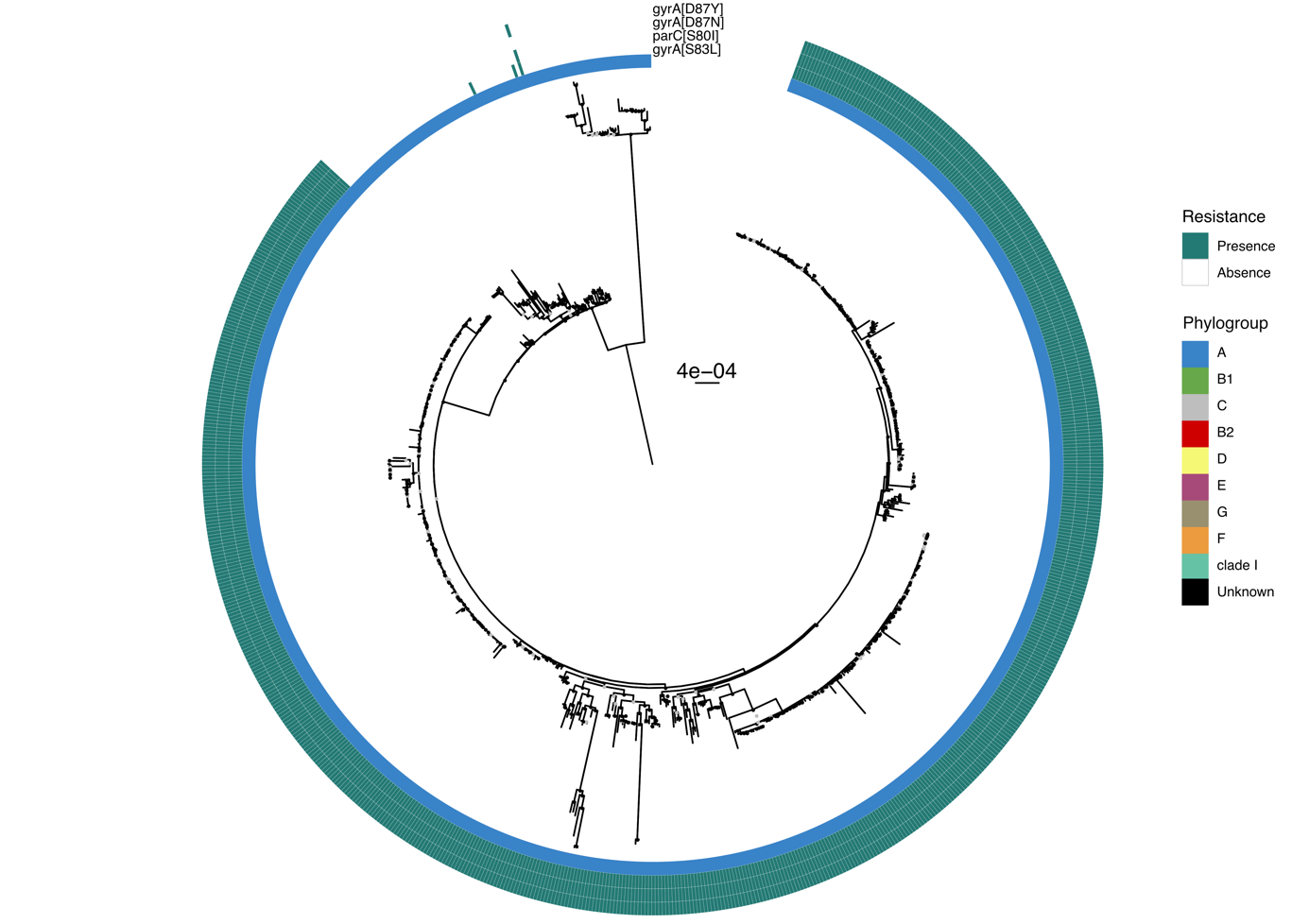

**Figure S5: Phylogeny and distribution of quinolone resistance mutations in DNA gyrase and topoisomerase IV genes for E. coli of the ST410 dataset.** The maximum likelihood phylogenetic tree of the species and the scale of substitution per site are in the center with IQTree (model GTR+F+I+G4 and 1000 ultrafast bootstraps). The E. coli phylogroups are represented by the colors in the inner center. The marine green squares indicate the presence of the respective mutation across the species. Ultrafast bootstrap values superior to 75% are shown with a light gray circle and values superior to 90% with a black circle.

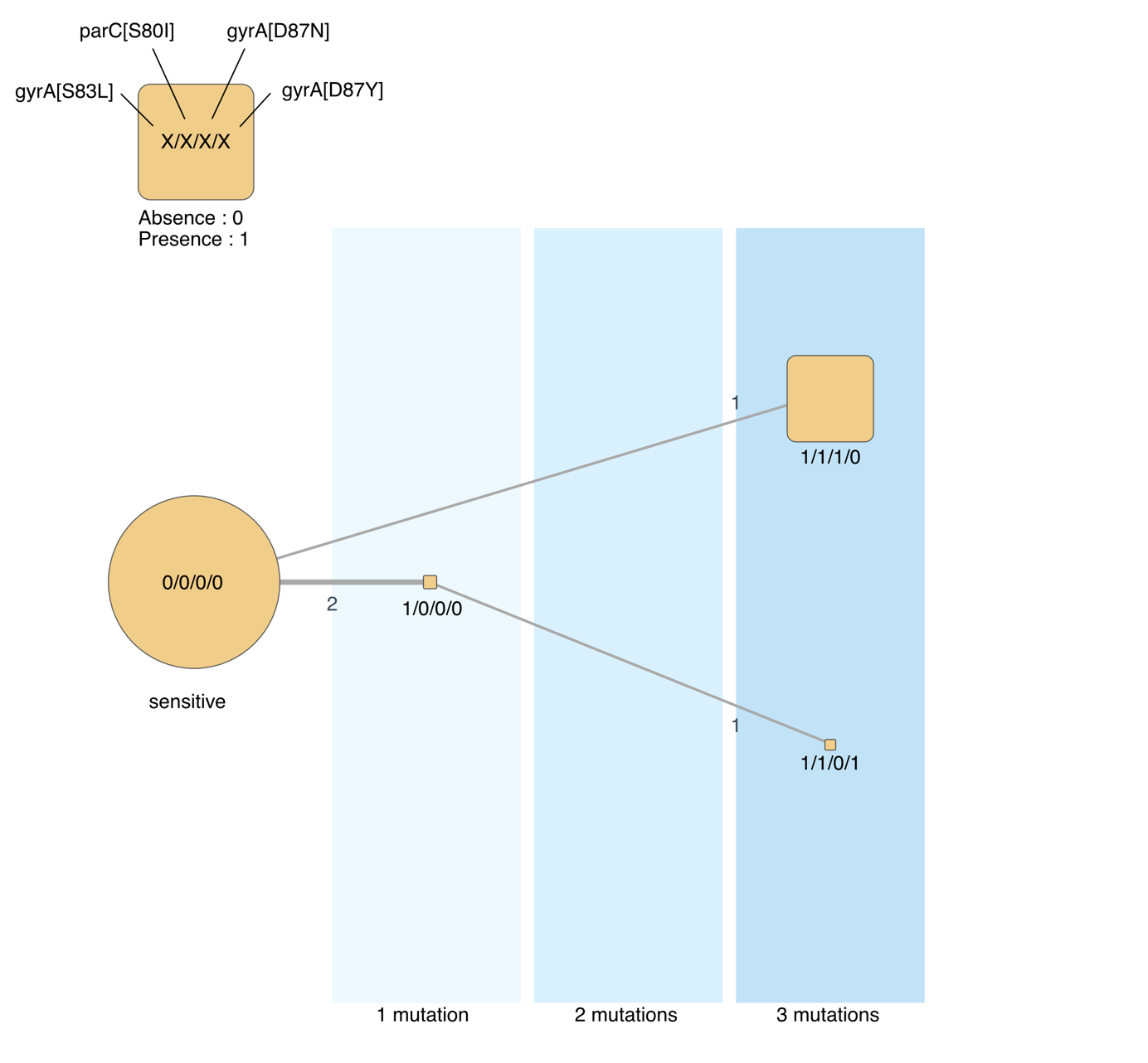
**Figure S6: Trajectories of acquisition of the main resistance mutations in E. coli for the ST410 dataset.** The blue areas represent the number of distinct mutations in genomes. The size of the square scales with the number of genomes observed (leaves of the tree). The edges represent the chronologies of acquisition of one or several mutations as inferred from the reconstruction of ancestral states on the tree. The edge size is proportional to the frequency of the respective transition, with the labels showing this exact number.

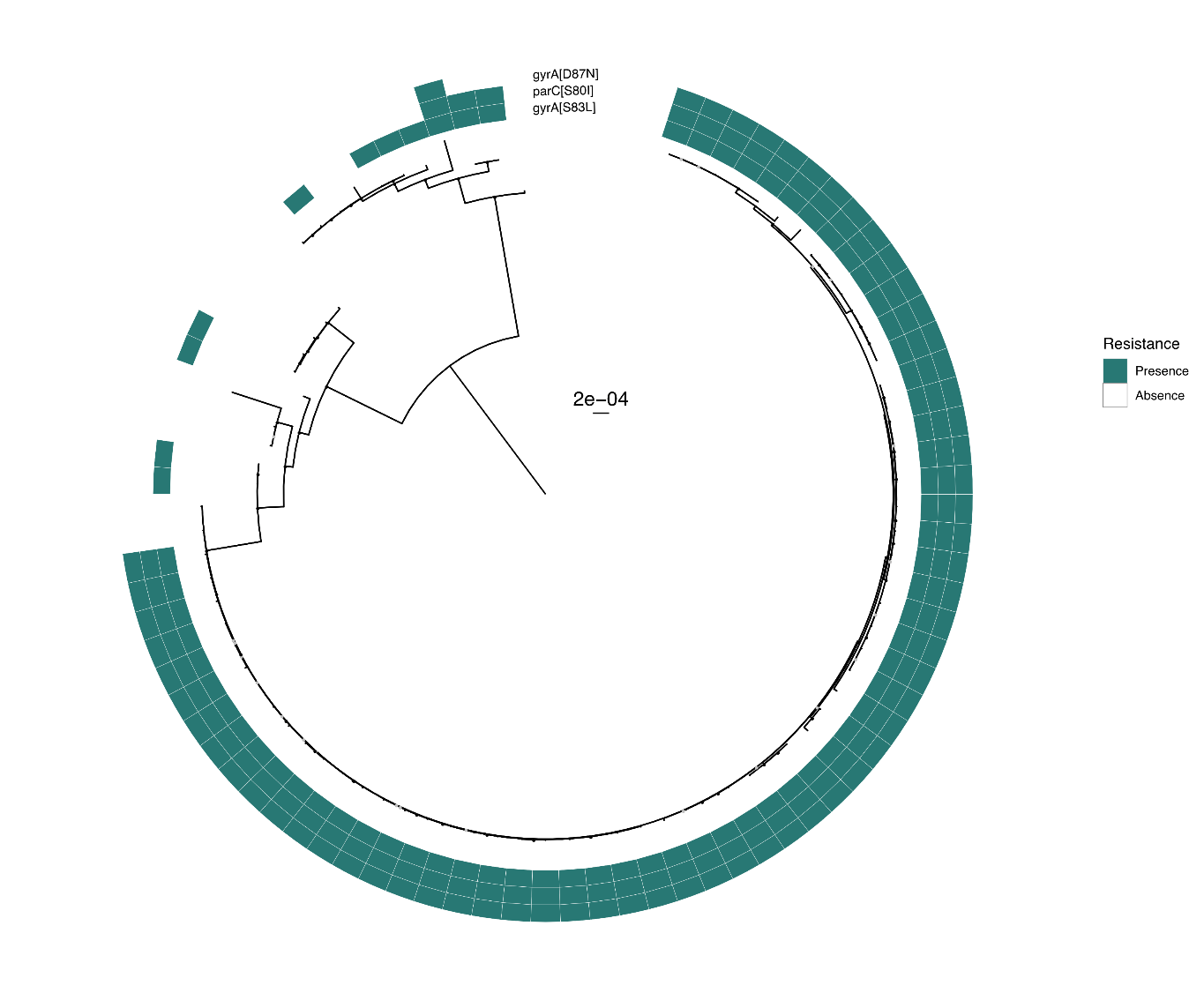

**Figure S7: Phylogeny and distribution of quinolone resistance mutations in DNA gyrase and topoisomerase IV genes for E. coli of the ST131 dataset.** The maximum likelihood phylogenetic tree of the species and the scale of substitution per site are in the center with IQTree (model GTR+F+I+G4 and 1000 ultrafast bootstraps). The marine green squares indicate the presence of the respective mutation across the species. Ultrafast bootstrap values superior to 75% are shown with a light gray circle and values superior to 90% with a black circle.

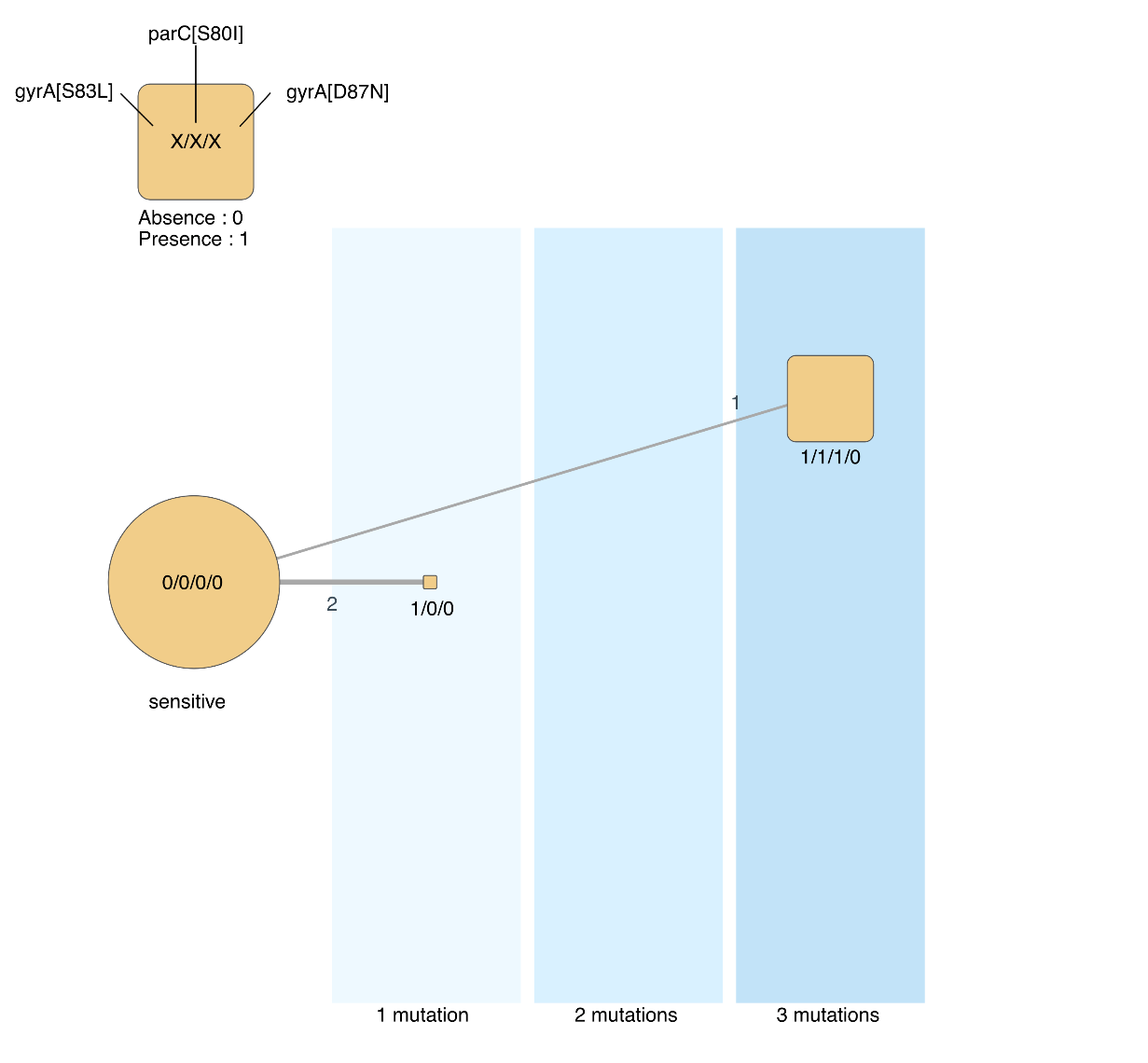

**Figure S8: Trajectories of acquisition of the main resistance mutations in E. coli for the ST131 dataset.** The blue areas represent the number of distinct mutations in genomes. The size of the square scales with the number of genomes observed (leaves of the tree). The edges represent the chronologies of acquisition of one or several mutations as inferred from the reconstruction of ancestral states on the tree. The edge size is proportional to the frequency of the respective transition, with the labels showing this exact number.

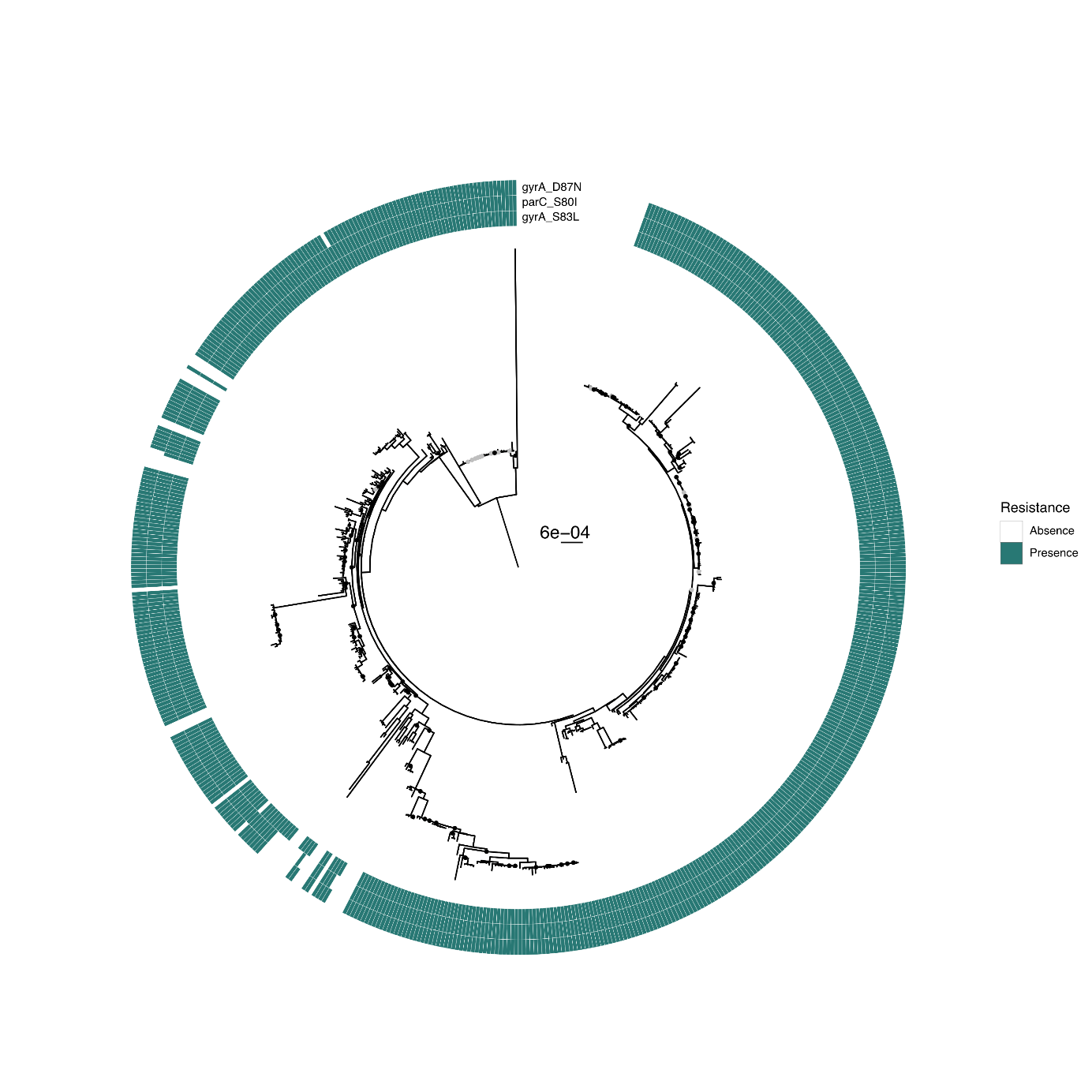

**Figure S9: Phylogeny and distribution of quinolone resistance mutations in DNA gyrase and topoisomerase IV genes for E. coli of the ST167 dataset.** The maximum likelihood phylogenetic tree of the species and the scale of substitution per site are in the center with IQTree (model GTR+F+I+G4 and 1000 ultrafast bootstraps). The marine green squares indicate the presence of the respective mutation across the species. Ultrafast bootstrap values superior to 75% are shown with a light gray circle and values superior to 90% with a black circle.

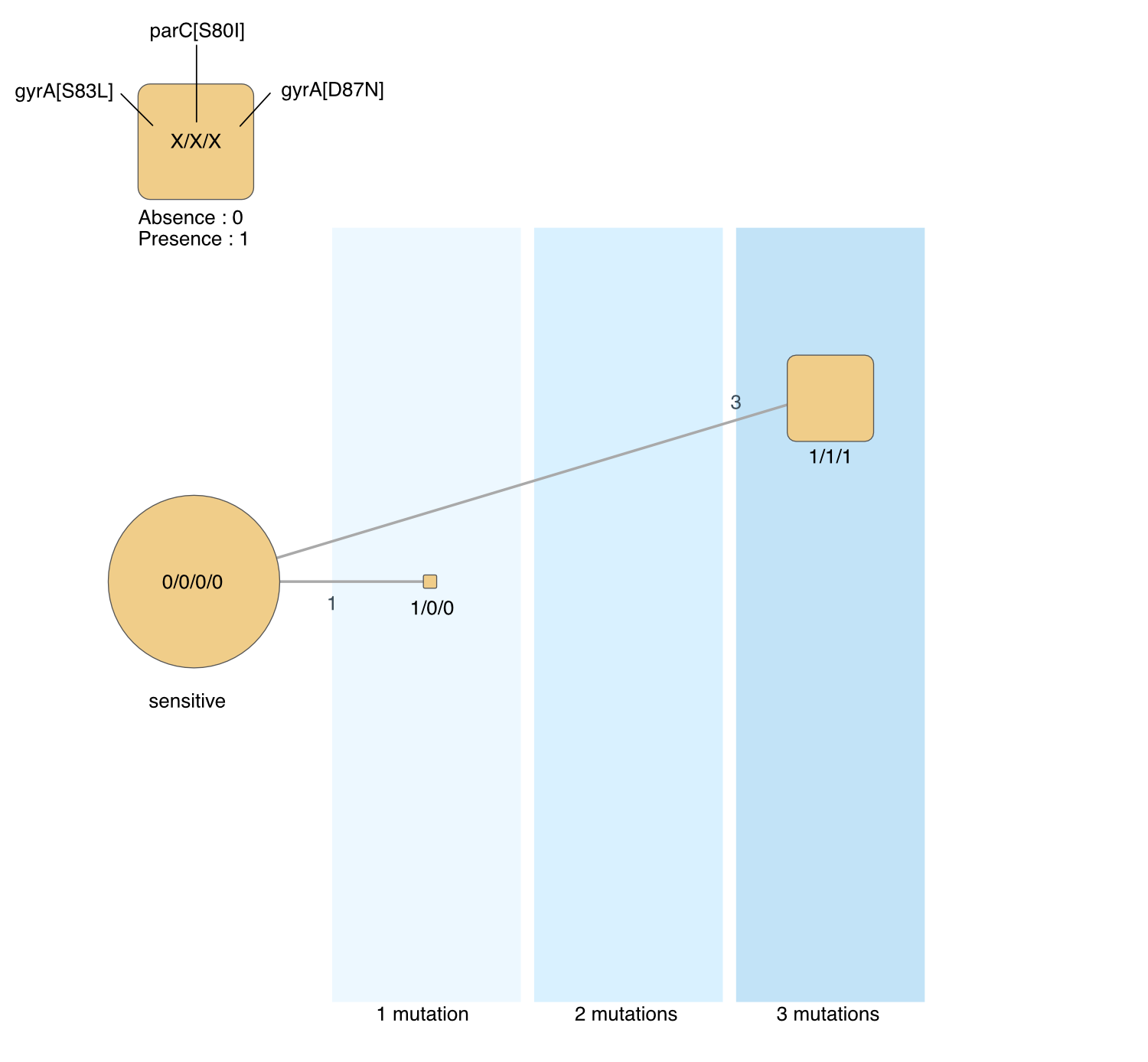

**Figure S10: Trajectories of acquisition of the main resistance mutations in E. coli for the ST167 dataset.** The blue areas represent the number of distinct mutations in genomes. The size of the square scales with the number of genomes observed (leaves of the tree). The edges represent the chronologies of acquisition of one or several mutations as inferred from the reconstruction of ancestral states on the tree. The edge size is proportional to the frequency of the respective transition, with the labels showing this exact number.

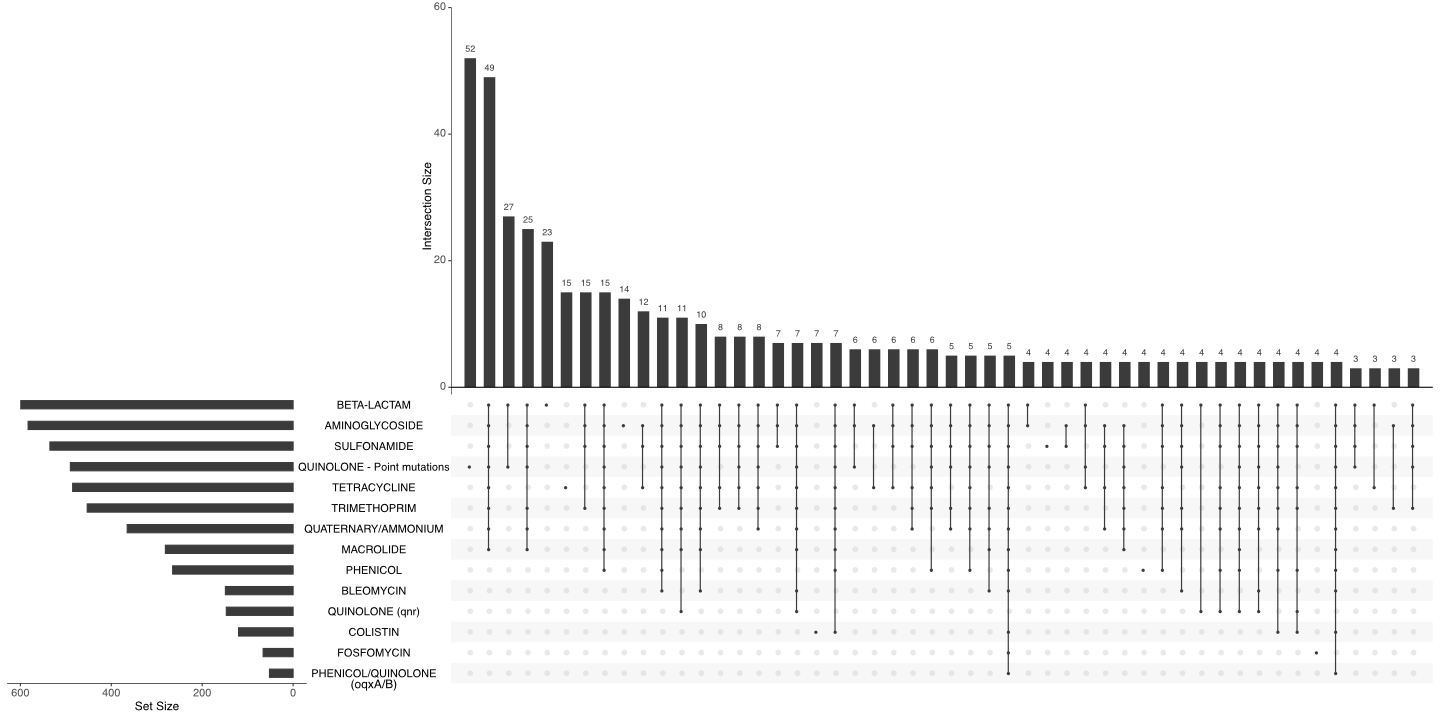

**Figure S11: Upset plot of the different AMR classes in the RefSeq dataset.** The horizontal bar plot represents the distribution of the mutations in the RefSeq dataset. The vertical bar plot represents the distribution of the AMR class combinations. Lines and dots represent the different combinations.

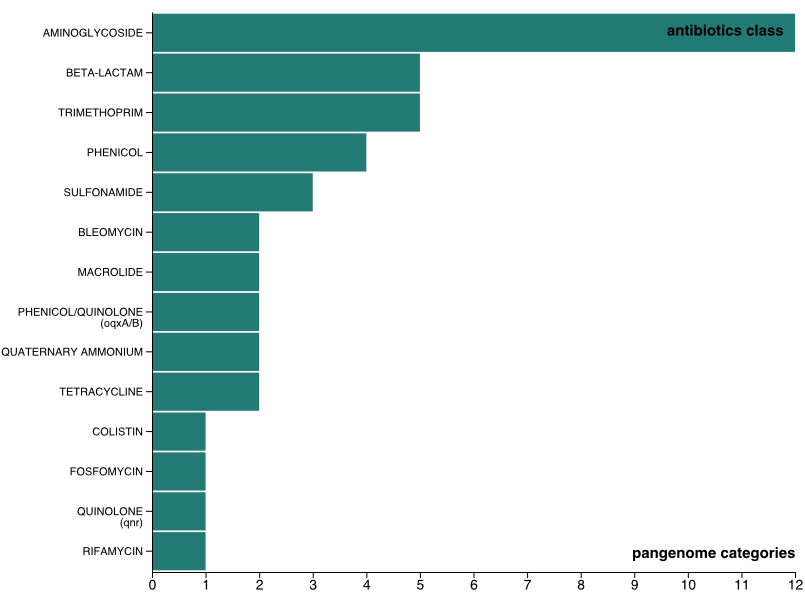

**Figure S12: Distribution of the different class of antibiotic frequently acquired after the acquisition of the quinolone conferring mutations in the RefSeq dataset.** Antibiotic classes are ordered according to the number of pangenome families acquired after the resistance mutations.
